# Atlas of adult Danionella fish telencephalon reveals retention of larval characters

**DOI:** 10.64898/2026.09.23.753900

**Authors:** Jonathan T. Perelmuter, Andrew H. Bass

## Abstract

The adult size of the vertebrate telencephalon typically impedes our interpreting its functions within the context of brain-wide circuit dynamics. *Danionella*, a genus of miniature and transparent teleost fishes closely related to zebrafish (*Danio rerio*), holds promise for whole-brain imaging and connectomics in an adult vertebrate. These advantages result from progenesis, a form of heterochrony characterized by early developmental onset of sexual maturation and cessation of growth leading to the retention of larval or juvenile characters in adults. Using molecular markers, tract tracing, and quantitative comparisons of cellular distribution patterns, we report a detailed neuroanatomical atlas of the adult *Danionella dracula* (“Dracula fish”) telencephalon. The results demonstrate that adults of Dracula fish and other *Danionella* species show extreme reduction in cellular migration from periventricular zones and topographic reorganization of major pallial divisions, including a compartment-like caudal pallium reminiscent of insect glomeruli (a dense neuropil surrounded by a thin layer of somata). We further document these characters in larval zebrafish, providing essential evidence corroborating a larval-like telencephalon in *Danionella* adults. Despite this phenotype, most canonical regions of the adult teleost telencephalon are identified. We also generate a transgenic GCaMP Dracula fish line, an annotated 3-dimensional atlas, and align functional imaging data to demonstrate the usefulness of this atlas as a resource for deciphering how the novel progenetic cytoarchitecture of the adult *Danionella* telencephalon can support its rich behavioral repertoire.

## INTRODUCTION

The successful performance of naturally selected behaviors is thought to depend upon neural activity across the telencephalon (e.g., including but not limited to amygdala, basal ganglia, cortex, hippocampus)^1^ and interconnections with the rest of the brain.^2^ A mechanistic understanding of telencephalic function demands measuring and manipulating the activity of individual neurons embedded in brain-wide circuits during behavior. In nearly all vertebrates, this ambition is thwarted by brain size and current technological limitations. A prominent exception has been optical interrogation of the brain in zebrafish (*Danio rerio*) at transparent, larval stages.^3,4^ Limitations with larvae include a telencephalon that is not fully developed,^5–8^ and the lack of many adult-typical behaviors.^9–14^ *Danionella* fishes, a recently described group of miniature fishes^15–19^ closely related to zebrafish,^17,20–22^ have begun to address these shortcomings.^23–36^ Adult, sexually mature *Danionella* are transparent, lack a skull roof (Figure 1A), and possess a brain volume of ∼0.6 mm^3^, one of the smallest adult vertebrate brains on record.^23^ In comparison, the adult mouse and zebrafish brains are estimated at 400-500 mm^3^ and 2.8 mm^3^, respectively.^37,38^ Coupled with amenability to genetic manipulations validated in zebrafish, the diminutive *Danionella* holds the promise for whole-brain connectomics and brain-wide optical recording and circuit perturbations at single-cell resolution during adult behaviors.^26,29,30,36,39–41^

**Figure 1.**
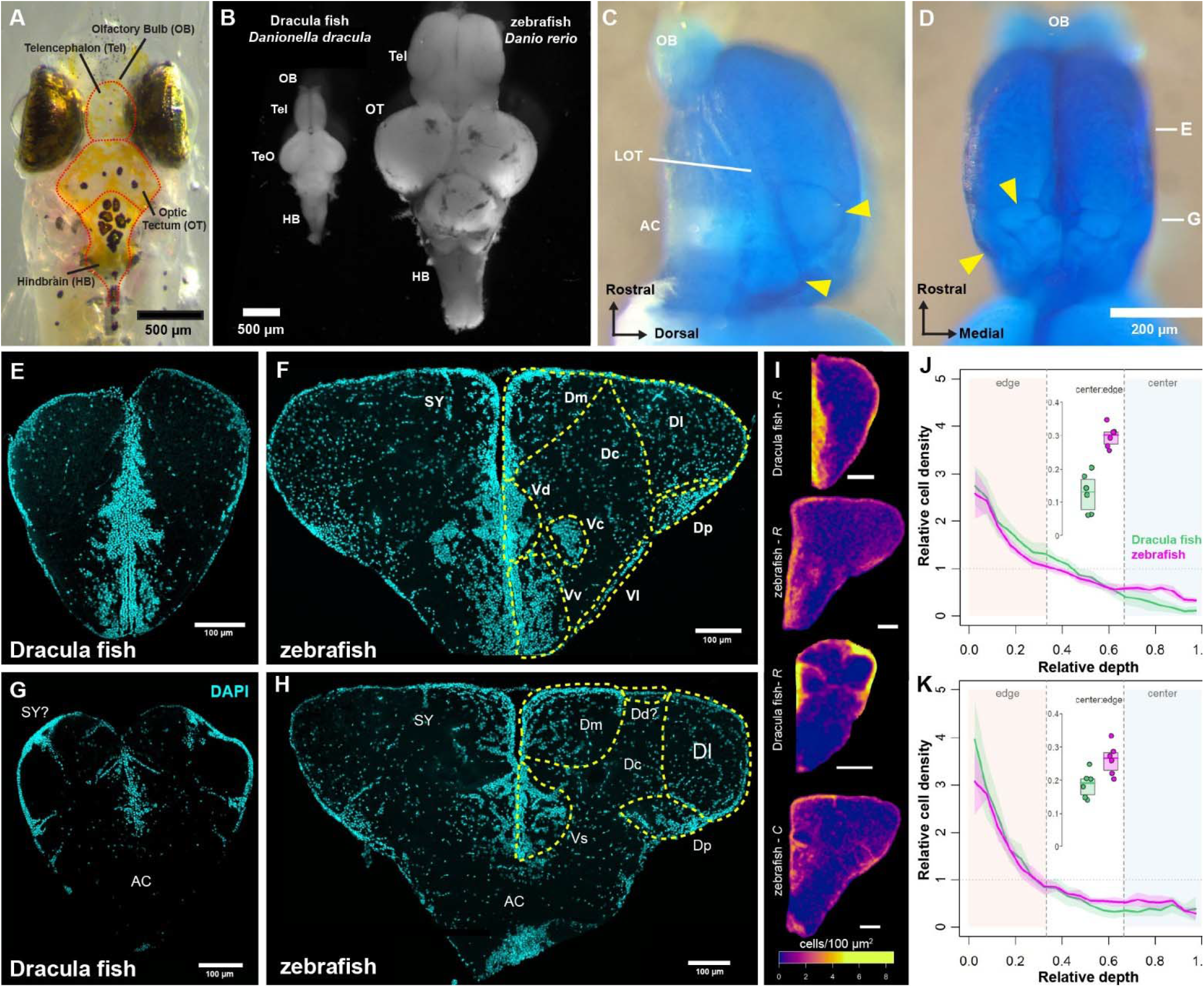
Reduced cellular migration in adult Dracula fish telencephalon as compared to adult zebrafish. (A) Dorsal view of Dracula fish head. (B) Adult Dracula fish and zebrafish brains. (C and D) Lateral (C) and dorsal (D) view of Dracula fish telencephalon and OB stained with methylene blue (AC, anterior commissure; LOT, lateral olfactory tract). Yellow arrowheads point to compartment-like caudal pallium. Letters indicate position of transverse sections in (E) and (G). (E-H) Comparison of rostral and caudal 12-µm transverse sections through telencephalon of adult Dracula fish (E, G) and zebrafish (F, H). DAPI (cyan) labels cell nuclei. For nomenclature and abbreviations, see Results section titled *Telencephalic nomenclature*. (I) Hemisection cell density heatmaps compare Dracula fish to zebrafish (bottom) at rostral (*R*) and caudal (*C*) levels (n = 6 per group). (J,K) Relative cell density plotted across depth (hemi-section edge to center) compares Dracula (green) and zebrafish (magenta) rostrally (J) and caudally (K), with 95% confidence bands (n = 6). Dotted horizontal line at 1 is density assuming a uniform distribution. Inset box plots show center-to-edge ratios. Lower ratio indicates a sparser interior relative to the hemi-section edge. Dracula has a lower ratio rostrally (p = 0.0022, Cliff’s delta = −1) and caudally (p = 0.026, Cliff’s delta = −0.78).

Telencephalic cytoarchitecture is widely variable across bony vertebrates and especially in ray-finned fishes,^1,42^ the radiation of bony vertebrates that includes *Danionella,* zebrafish, and more than half of all living vertebrate species.^43,44^ This variation is especially true for the pallium, the dorsal division of the telencephalon that undergoes a unique developmental process in actinopterygians termed eversion. Eversion alters the adult positions of pallial divisions relative to those in the two other major vertebrate radiations, cartilaginous and lobe-finned fishes, which includes tetrapods.^45,46^ The degree of rearrangement, still debated, appears to vary across ray-finned fishes, further complicating inter-species comparisons.^47^

Crucially, what makes *Danionella* potentially transformative for developmental biology and neuroscience likely results from progenetic paedomorphosis, a form of development in which the onset of sexual maturation speeds up relative to the body.^48,49^ The result is an adult that retains larval and juvenile characters of its ancestors. *Danionella dracula* (Dracula fish), our focal study species, sits at the base of the genus phylogeny, and lacks nearly one-third of the bones found in adult zebrafish and most cyprinids.^17,50^ Together with life-long transparency and lack of scales that make the entire brain optically accessible,^25,27–30^ progenesis in Dracula fish and other *Danionella* species appears to manifest as an organism-wide developmental truncation.^17,50,51^ It remains unknown if this profound truncation extends to the telencephalon. Does it retain larval characters despite displaying behaviors that typically emerge later in development (social affiliation,^23,24^ associative learning,^32,52^ sound production, courtship, and aggression^23,25,53,54^)?

Here, we report that the telencephalon in Dracula fish retains larval-like characters. Hence, despite their close phylogenetic relationship, a simple transposition of an adult zebrafish brain atlas^38,55^ to *Danionella* is not possible. To generate an atlas with sufficient detail to guide experimental investigations and informed comparisons to other fishes and vertebrates in general, we document for adults the expression of calcium-binding proteins, neuromodulators, transcription factors and olfactory-bulb projections. Since most information for zebrafish is limited to early larval or adult stages,^5,38,55–57^ we examine their telencephalon at 14-28 days-post-fertilization (dpf), identifying characters in Dracula fish that map onto larval and adult stages in zebrafish. This helps corroborate the retention of larval characters in adults. Together, the results enable us to identify most of the major divisions of the telencephalon of ray-finned fishes in Dracula fish. Comparison with two other *Danionella* species (*D. priapus* and *D. cerebrum*) confirms a genus-wide signature of telencephalic paedomorphosis.

This atlas of the telencephalon will prove an essential tool for researchers currently studying or considering adoption of *Danionella* fishes as a model clade^58,59^ for investigating brain-wide circuitry enabling adult vertebrate behaviors.^24^ More broadly, this atlas enhances the ability to integrate future functional imaging, molecular profiling and connectomics for *Danionella* fishes with the wealth of knowledge accrued from zebrafish and other vertebrates, allowing the extraction of general principles underlying the evolution and development of vertebrate brain structure-function relationships.

## RESULTS

The adult Dracula fish brain superficially resembled an adult zebrafish brain (Figures 1A and 1B). Closer examination of the telencephalic surface revealed a unique cytoarchitectural pattern made more salient after incubation in methylene blue to visualize cell bodies. Rostral to the anterior commissure (AC), cells were uniformly distributed across the surface, while at the AC and more caudal, a series of cell-dense bands gave the pallium a lobe-like appearance (arrowheads, Figures 1C and 1D; see also Figure 3 in^27^).

### Cell migration appears reduced in adult Dracula fish telencephalon

To compare telencephalic cytoarchitecture between adult Dracula fish and adult zebrafish, we examined transverse sections labeled with DAPI (Figures 1E-1H). Both species showed tightly packed cell nuclei along the midline ventricle surface and dorsomedial ventricular surface (due to eversion^45,46,60–63)^. However, overall cellular density in Dracula fish (Figures 1E and 1G) decreased substantially distal to the ventricle, unlike adult zebrafish which exhibited substantial migration (Figure 1F and 1H). This decrease was most extreme in Dracula fish caudal pallium (Figures 1G, S1A and S1B). Dense bands of cells framed deeper, cell-sparse compartment-like regions, hence the impression of lobes in surface views (Figures 1C and 1D).

Immunolabeling of neurons with Hu (Figures S1C and S1D) and synaptic vesicle protein 2 antibodies (SV2, Figures S1E and S1F) confirmed these patterns and suggested a dense, synaptically-rich neuropil throughout the brain.

To quantify species difference in cytoarchitectural patterns, we segmented cell nuclei in hemisections at rostral and caudal levels of adult Dracula and zebrafish telencephalon (n = 6 per species and level). We extracted cell centroids, warped sections within each group into a common reference frame and generated averaged heatmaps of cell density (Figure 1I). While relative cell density was highest at the midline and peripheral edges and declined toward the center in both species, this decline was steeper in Dracula fish (Figures 1J, 1K and S1G, Rostral: −2.53 vs. −1.93 median slope, δ = −0.89, p = 0.0087, Caudal: −2.69 vs −2.24 median slope, δ = −0.83, p = 0.0152, Mann-Whitney U tests). To summarise this decline in a single size- and shape-independent value, we computed the center:edge ratio (Figure S1H) for each animal, where smaller values indicate a cell-sparse interior relative to the periphery. Dracula fish had a lower ratio than zebrafish rostrally (0.13 vs. 0.30 median ratio, δ = −1, p = 0.0022) and caudally (0.19 vs 0.26 median ratio, Cliff’s δ = - 0.78, p = 0.026, Mann-Whitney U tests).

*Telencephalic nomenclature*. Parcellation of the ray-finned fish telencephalon traditionally follows a topographic nomenclature originated by Nieuwenhuys,^45,64,65^ with modifications to reflect new empirical evidence, varying interpretations and interspecific differences.^61,62,66^ The pallium, or area dorsalis (D), generally comprises five major divisions: medial (Dm), dorsal (Dd), lateral (Dl), posterior (Dp) and central (Dc) (see Figures 1F and 1H). The definition and boundaries of Dd and Dc in zebrafish remain disputed.^67–69^ The subpallium, or area ventralis (V), comprises four nuclei rostral to the AC: dorsal (Vd), ventral (Vv), lateral (Vl) and central (Vc). Starting at the AC, Vd is replaced caudally by supracommissural (Vs), and then postcommissural (Vp) and intermediate (Vi) nuclei. The subpallium also includes dorsal (ENd) and ventral (ENv) entopeduncular nuclei. Most of this map is supported by molecular markers and, more recently, single-cell transcriptomics.^70–72^ However, the paucity of cells distal from the periventricular zones confounded discernable cytoarchitectural patterns characteristic of zebrafish and other teleosts, hindering our ability to readily identify these canonical regions.

### Distinguishing pallial (D) - subpallial (V) boundary

Parcellation of telencephalon into pallium and subpallium is fundamental to all vertebrates and occurs early in embryogenesis.^1,73^ Despite the lack of cytoarchitectural differentiation in Dracula fish, we reasoned the D-V boundary should be identifiable using evolutionarily conserved marker genes and proteins.

#### Calretinin suggests D-V boundary

Immunostaining for calretinin, a calcium binding protein distinguishing the pallium from the subpallium in ray-finned fishes,^69,74–78^ provided initial evidence for the D-V boundary. A sagittal view revealed a medial, ventrally located cluster of calretinin positive (+) somata extending as an arc across the rostro-caudal extent of the telencephalon (arrows, Figure 2A). Transverse sections showed a dorsally located triangular-shaped field of labeled processes (henceforth referred to as a “field”), centered around a wedge of somata along the midline ventricle (yellow arrowheads, Figures 2B-2D). In sagittal view, this calretinin field arced across the telencephalon (yellow arrowheads, Figure 2Ai) and became especially dense past the AC (white asterisks, Figures 2A and 2Ai). We designated this area Dm, since this calretinin field in zebrafish identifies Dm^78^, which sits at the ventral most extent of the medial pallium (Figures 1F and 1H).

**Figure 2.**
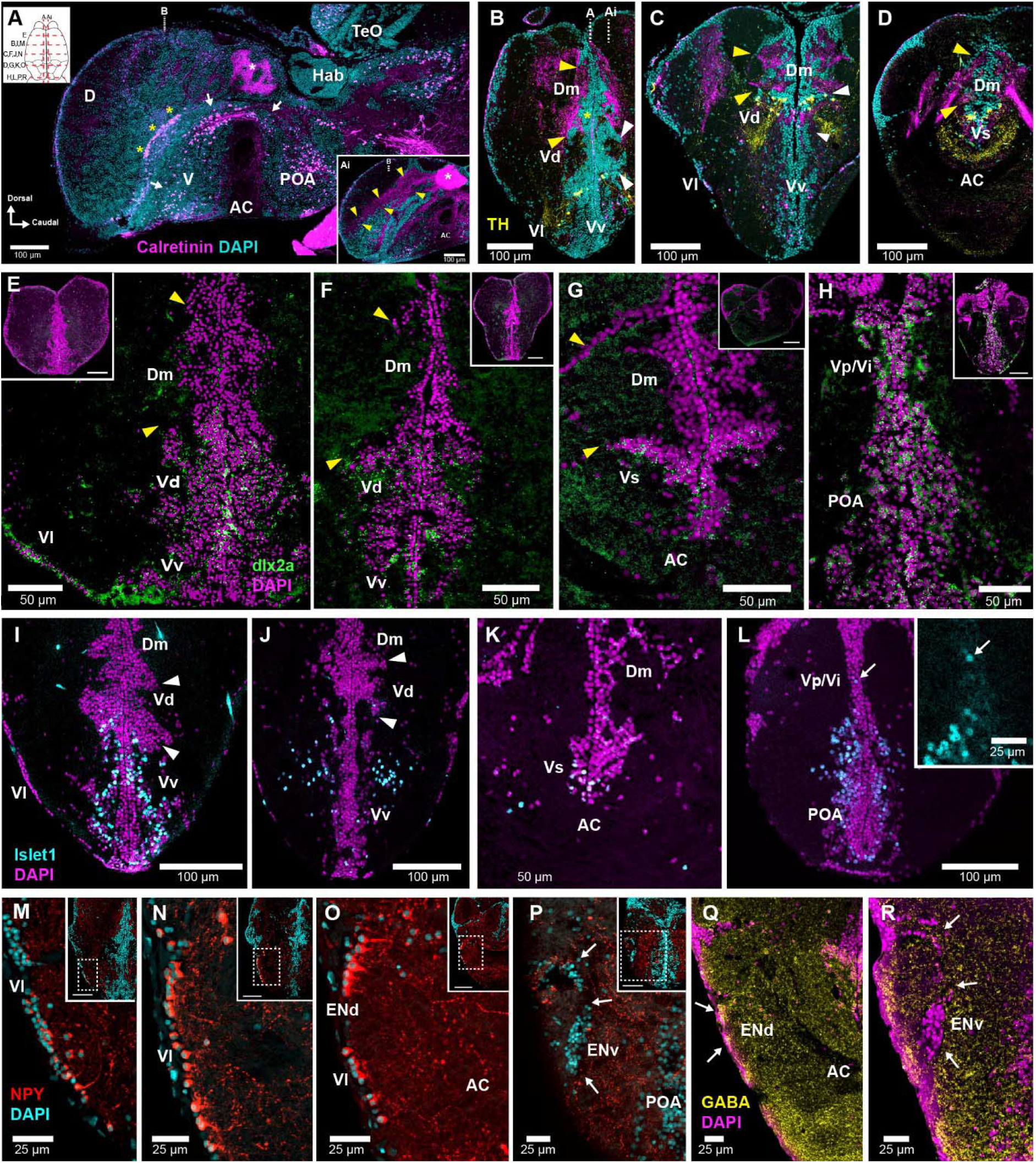
Identifying the boundary and subdivisions of pallium (D) and subpallium (V) in Dracula fish. (A, Ai) Sagittal sections showing calretinin(magenta) and DAPI (cyan). White arrows (A) show calretinin+ cells arc over anterior commissure (AC) and preoptic area (POA). Dorsal schematic indicates levels of sections in (A-R). Hab = habenula; TeO = optic tectum. (B-D) Tranverse sections, rostral to caudal, with calretinin and tyrosine hydroxylase (TH, yellow). Yellow arrowheads (Ai, B-D) indicate calretinin+ fibers within Dm. Dotted white lines in (B) indicate position of sagittal sections (A, Ai). Yellow asterisks track midline ventricle-contacting calretinin+ cells (see Figure S2A). (E-H) HCR-ISH for *dlx2a* (green) and DAPI (magenta) in transverse sections. Yellow arrowheads correspond to calretinin+ Dm fibers in (B-D). Scale bars = 100 µm in lower magnification insets. (I-L) *Islet1* (cyan) and DAPI (magenta) locates Vd, Vv, and Vs. White arrowheads (B,C,I,J) denote dorsal and ventral extent of Vd. Arrows (L) indicate sparse *islet1* labeling in Vp/Vi compared to POA. (M-P) Neuropeptide Y (NPY, red) and DAPI (cyan) distinguishes Vl and ENd. Arrows in (P) show ENv null for NPY. Scale bars in insets = 100 µm. (Q and R) ENd has GABA+ (yellow) cells (Q), while ENv is null for GABA (R).

#### TH identifies V subnuclei

To confirm the D-V boundary and identify Vd and Vv, we immunolabeled sections for calretinin and tyrosine hydroxylase (TH, indicative of catecholamine synthesis).^79^ A cluster of TH+ somata appeared just caudal to the olfactory bulbs, associated with two, teardrop-shaped midline-positioned cell clusters just under the D-V boundary defined by calretinin, which we identified as Vd (Figure 2B, white arrowheads). TH+ somata continued caudally under Dm’s calretinin+ field (Figure 2C). We identified Vs by a midline-positioned cluster of TH+ somata lying above a dense band of processes extending across the AC (Figure 2D). Caudal to the AC, there were no subpallial TH+ somata, although some appeared in the ventral POA (Figure S2B). A similar pattern of TH+ somata largely associated with Vd, and also dopaminergic, is in zebrafish and considered a conserved feature of the teleost telencephalon.^80–82^

#### Dlx2a corroborates D-V boundary

*Dlx*-family homeobox genes specify GABA neuron differentiation across the vertebrate forebrain, with expression that persists into adulthood throughout the subpallium.^83–87^ Rostral to the AC, hybridization chain reaction fluorescence *in situ* hybridization (HCR-FISH) showed *dlx2a* (teleost ortholog to *Dlx2*) expression throughout the ventral telencephalon (V, Figures 2E and 2F). In agreement with the boundary suggested by calretinin, *dlx2a* expression extended to the ventral border of Dm (lower yellow arrowheads, Figures 2B, 2C, 2E, 2F). The dorsal extent of *dlx2a* matched a small cluster of calretinin+ somata directly contacting the ventricle, which followed the aforementioned arc of ventral calretinin somata (yellow asterisks and arrows, respectively, Figures 2A and 2B). This midline cluster marks the precise boundary between pallium and subpallium (Figure S2A), offering a potentially useful landmark when viewing the telencephalon in the horizontal plane (e.g., during calcium imaging). A thin strip of *dlx2a*+ cells ran along the ventrolateral edges of the subpallium which we labeled Vl (Figure 2C), consistent with its location in other teleosts (Figure 1F).^45,62^ At commissural levels, a small cluster of *dlx2a+* cells appeared above the AC, identifying Vs (Figure 2G). Posterior to the AC, *dlx2a+* expression was extensive throughout the midline, including the preoptic area (POA) and Vp/Vi (Figure 2H). Immunostaining for GABA closely matched *dlx2a* expression, showing strong labeling throughout V, with the dorsal boundary apposing Dm (Figure S2C, compare to 2E and 2F).

#### Islet1, NPY identify subpallial nuclei

In zebrafish adults, *islet1*, a highly conserved transcription factor expressed in the subpallium,^88,89^ identifies Vv.^86,87,90^ While absent from most of Vd, it has been reported in a ventral division of Vd.^90^ In Dracula fish, immunohistochemical labeling for *islet1* was restricted to a ventral area around the midline (Figure 2I and 2J). Expression was largely absent from Vd, although some *islet1*+ somata appeared consistently in Vd’s ventral aspect (Figures 2I and 2J). Although absent from Vl, *islet1*+ somata occurred in Vs (Figure 2K) and throughout the post-commissural ventral midline, where a higher concentration of *islet1*+ cells distinguished the POA from near absence of labeling in the more dorsal Vp/Vi (Figure 2L). In zebrafish adults, *islet1* is expressed partly in Vs, not at all in Vp, with few *islet1+* cells in Vi.^90–92^

Neuropeptide Y (NPY) immunohistochemistry confirms Vl and helps distinguish it from ENd in adult zebrafish.^71,78^ In Dracula fish, NPY+ somata formed a thin strip along the ventrolateral edge of the subpallium, corresponding to *dlx2a+* and GABA+ Vl cells (Figures 2E and S2C), extending from caudal to the olfactory bulbs almost to habenula (Figures 2M-2O). Around the AC, NPY+ somata at the dorsal pole of this strip were larger with more discernable processes, while smaller ventral NPY+ cells disappeared at more caudal levels (Figure 2O). We label this dorsal division the ENd. While NPY+ cells are found in zebrafish Vd, we saw none in Dracula fish. However, we consistently found a few scattered NPY+ somata between Vl/ENd and Vd/Vv (arrows, Figure S2D). A small band of NPY-somata replaced ENd at the caudal end of the telencephalon, which we identified as ENv (arrows, Figure 2P). Like adult zebrafish,^78,93^ ENd is NPY+ and GABAergic (Figures 2O and 2Q), while ENv is null for both markers (Figures 2P and 2R). Finally, there was no candidate cell group resembling the Vc in zebrafish (see Figure 1F) and other teleosts.^55,94,95^

#### Olfactory bulb (OB) connectivity, TH, otpa confirm identifications

OB connectivity to the telencephalon, extensively studied for teleosts,^62,69,76,96–98^ along with TH+ immunohistochemistry aided identification of telencephalic nuclei. Small, unilateral OB injections of neurobiotin (Figure 3A, inset) resulted in dense retrograde filling of somata and processes in Vd and Vv, with the most in Vv (Figure 3A’); TH+ label corroborated a Vd-Vv boundary (Figure 3A’’). Consistent with anterograde label,^99,100^ neurobiotin-filled, terminal-like puncta appeared over lightly-stained somata at the lateral edge of D from mid-rostral until AC levels (arrows, Figure 3B). The most prominent labeling of puncta and processes within the pallium occurred in its ventral and caudal-most compartment (Figures 3C-3E), confirming it as Dp, the main olfactory recipient pallial division of teleosts.^62^ Neurobiotin label in Dp extended past the AC to a dense “cap” of cells at the caudal end of the telencephalon, many of which exhibited retrograde labeling (Figures 3D and 3E). Consistent with studies in zebrafish,^96^ OB input was greater in the right dorsal habenula (Figure 3E). Dp also possessed dense TH+ label at rostral levels above the AC (Figure 3F). OB-recipient Vs and Vi had sparse populations of densely-filled neurobiotin+ somata and puncta (Figures 3C and 3D).

**Figure 3.**
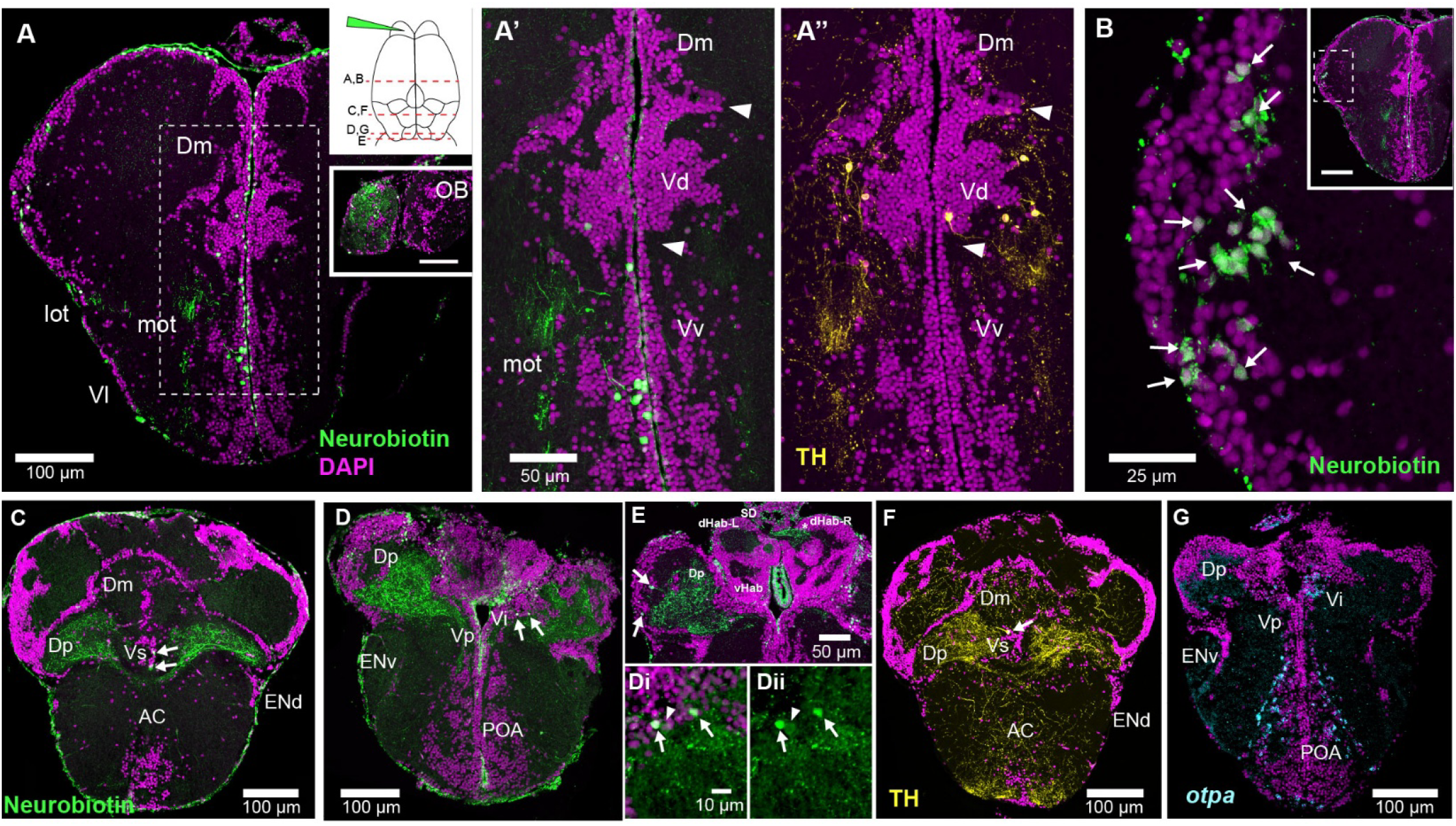
Olfactory bulb efferents, TH and *otpa* confirm divisions of pallium (D) and subpallium (V) in Dracula fish. (A-G) Transverse sections; inset dorsal schematic (A) indicates levels. (A) Neurobiotin labeled (+, green) fibers and cells following injection into OB (inset). Closeups show (A’) filled cells concentrated in Vv and (A’’) TH+ cells in Vd. Arrowheads mark extent of Vd. (B) Anterograde-filled cells at lateral edge of D. Arrows indicate filled cells; note neurobiotin+ puncta over fainter neurobiotin signal in cells. (C-E) Neurobiotin+ fibers and cells (arrows) in Dp, Vs and Vi. Closeup of neurobiotin+ cells in Vi shown with (Di) and without (Dii) DAPI. Arrowhead shows fainter neurobiotin+ cell. (E) Asterisk indicates denser neurobiotin+ fibers in dorsal right habenula (dHab-R). Abbreviations: dHab-L = dorsal left habenula; SD = saccus dorsalis; vHab = ventral habenula. (F) TH+ fibers in Dp. Arrow indicates TH+ cell in Vs. (G) HCR-FISH shows *otpa*+ cells in POA and Vi. Note absence of *otpa* in Vp. Cell nuclei labeled with DAPI (magenta). Scale bars in lower magnification insets = 100 µm.

Zebrafish Vi, recently proposed as a homologue to the mammalian medial amygdala,^101,102^ is distinguished from adjacent Vp by the transcription factor *otpa*.^92,102^ In Dracula fish, *otpa* expression was in the same subpallial region as neurobiotin fills, consistent with zebrafish Vi (Figure 3G). Like zebrafish,^92^ lack of *otpa* expression in Vp distinguished it from the *otpa*+ POA (Figure 3G).

### Topography of *Danionella* caudal pallium diverges from zebrafish

Two other *Danionella* species, *D. cerebrum* and *D. priapus*, revealed a caudal pallial cytoarchitecture of cell-sparse compartments like that of Dracula fish, suggesting a genus-wide character distinctly different from zebrafish (Figures 4A-4D). For all species, Dp was identified ventrolaterally. To confidently identify the topography of Dm, Dl and Dc in *Danionella* we compared calretinin and parvalbumin immunohistochemistry between the adults of zebrafish, Dracula fish, *D. cerebrum* and *D. priapus*.

**Figure 4.**
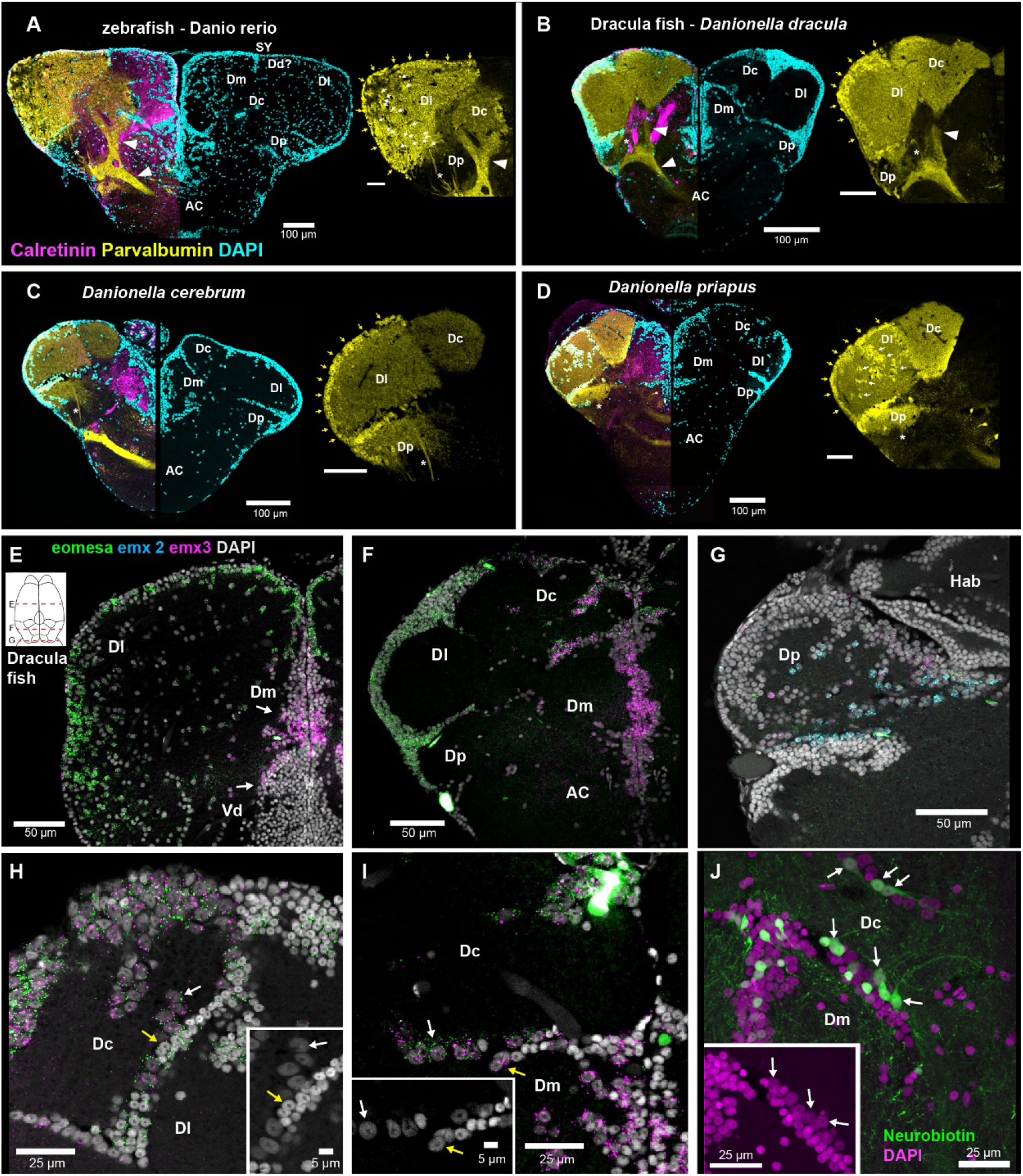
Comparison of adult *Danionella* species and adult zebrafish reveals topographic divergence of caudal pallium, confirmed with transcription factor expression. (A-D) Transverse sections at comparable caudal locations showing calretinin (magenta) and parvalbumin (yellow) in adult (A) zebrafish, (B) Dracula fish, (C) *Danionella cerebrum*, and (D) *Danionella priapus*. Right hemi-sections show only DAPI (cyan) to highlight cytoarchitectural patterns. Enlargement of parvalbumin labeling in pallium shown to right. Yellow arrows indicate parvalbumin+ cells at ventricular surface. White arrows indicate centrally migrated parvalbumin+ cells. Arrowheads in (A, B) denote parvalbumin fiber tracts between Dc and AC. Asterisks denote parvalbumin+ fibers entering AC that likely emanate from cells in Dl. (E-I) Transverse sections in Dracula fish with HCR-FISH showing *emx3* (magenta) in Dm, *eomesa* (green) in Dl, and both in Dc. Arrows in (E) indicate ventral region of Dm with greater *emx3* signal. (G) Caudal Dp shows *emx2* (blue) and *emx3* expression with sporadic *eomesa*. Dorsal view schematic (E) indicates approximate levels. DAPI (grey). (H-I) Cells expressing both *emx3* and *eomesa* are associated with Dc and have larger nuclei than in adjacent Dl (*eomesa+* only, H) and Dm (*emx3+* only, I). Insets DAPI only. White and yellow arrows indicate representative Dc and Dm/Dl cells, respectively. (J) Neurobiotin (green) filled cells in Dc and Dm. Inset shows Dm/Dc border with only DAPI (magenta). Filled cells projecting into Dc have larger nuclei (white arrows). See also Figure S3H. (H-J) at comparable position to (F, B).

*Dm.* A calretinin+ Dm in zebrafish extends from the subpallial border to the dorsal brain surface, bounded laterally by a parvalbumin+ filled Dl (Figure 4A).^69,78,103^ In Dracula fish, calretinin+ label did not reach the dorsal surface (Figure 4B), unlike *D. cerebrum* and *D. priapus*, which had a calretinin+ Dm location comparable to zebrafish (Figures 4A-4D). This suggested a uniquely ventral position for caudal Dm in Dracula fish.

*Dl*. As previously reported ^103,104^, caudal Dl in zebrafish was filled with parvalbumin+ fibers and somata, while a parvalbumin+ caudal Dc lacked labeled somata (Figures 4A and S3A). Parvalbumin+ processes and somata in all *Danionella* species’ most lateral pallial division appeared to correspond to caudal Dl in zebrafish. In Dracula fish and *D. cerebrum*, parvalbumin+ somata were largely periventricular and in two dense centrally extending bands forming caudal Dl’s borders (Figures 4B and 4C); *D. priapus* had clusters of parvalbumin+ somata throughout caudal Dl (Figure 4D). In contrast, parvalbumin+ somata in zebrafish were diffusely distributed across caudal Dl (Figures 4A and S3A). The processes of parvalbumin+ somata from the Dl/Dp border projected ventrally through Dp and then medially into the AC in all three Danionella species (Figures 4A-4D). Past the AC, Dp was lightly labeled compared to caudal Dl (Figure S3B) *Dc*. In Dracula fish, an expansive dorsal field of parvalbumin+ label, devoid of parvalbumin+ somata, directly overlaid Dm and extended ventrally toward the AC (arrowheads, Figure 4B). The comparable region in *D. cerebrum* and *D. priapus* was between Dm and Dl (Figures 4C and 4D). Despite its dorsal location in *Danionella* species, this region appears to correspond to caudal Dc in zebrafish.

### Transcription factor expression supports pallial identities in Dracula fish

To corroborate our identifications, we examined expression of transcription factors involved in forebrain patterning; *emx3*, *emx2 and eomesa* (also called *tbr2*).^105,106^ In adult zebrafish, *emx3* is strongly expressed throughout Dm and *eomesa* across most of Dl, while most of Dc expresses both.^107^ In Dracula fish, *eomesa* and *emx3* were largely non-overlapping rostral to AC, supporting delineation of the *eomesa*+ regions as Dl. *Emx3* expression matched the calretinin+ Dm (Figures 4E and 4F). Up to and including AC levels, *emx3* was stronger in ventral Dm (arrows, Figure 4E). Further caudal, *eomesa* and *emx3* were co-expressed throughout Dc (Figures 4F). Like adult zebrafish, Dracula fish Dc showed weak NPY and GAD expression compared to other pallial divisions^78^ (Figure S3C and S3D) and Dp somata were *eomesa*+ (Figure 4F).^107,108^ Caudally, Dp contained *emx3+* and sparse *eomesa+* cells (Figures 4G and S3E). It has been reported that *emx2* marks the caudal-most portion of Dc in adult zebrafish.^107^ However in Dracula fish, pallial *emx2* expression was confined to caudal Dp (Figure 4G and S3E). Dracula fish was *emx2+/ emx3+* in Vv (Figure S3F) and *eomesa*+ in ENv (Figure S3G).

In caudal pallium*, eomesa+*/ *emx3-* Dl cells and *emx3+/eomesa+* Dc cells intermingled within the cell dense borders between their neuropils (Figure 4H). Consistent with Dc containing larger cells than adjacent divisions in zebrafish and other teleosts,^62,69,76^ the DAPI-stained nuclei of *emx3+/eomesa+* Dc cells were consistently larger than adjacent *eomesa+*/ *emx3-* Dl cells. A similar pattern was observed at the Dc-Dm border (Figure 4I). Neurobiotin injections into rostral pallium (OB misses) showed the processes of smaller nucleated cells at the Dm-Dc border extending into Dm and larger cells into Dc (Figures 4J and S3H).

### *Danionella* adults and 14-dpf zebrafish larvae share Dc location

As Mueller et al.^104^ showed in zebrafish, parvalbumin+ fibers identifying caudal Dc continue rostrally where they ascend to the dorsal edge of D (Figure S3A). Although Dc was originally defined by its central position^45^, Mueller et al.^104^ proposed that it includes this rostral dorsal region. They also hypothesized that earlier in development, all of Dc occupies this dorsal position, but Dl and Dm subsequently grow to cover Dc at caudal levels. If correct, Dc’s dorsal position in adult *Danionella* would be paedomorphic. To investigate, we examined zebrafish larvae. Like adults (Figures 4A and S3A), two adjacent parvalbumin+ fields occurred near or at the AC at 14-, 21- and 28-days post fertilization (dpf): Dl - an intensely labeled lateral division with parvalbumin+ somata, and Dc - a medially adjacent region with lighter label, lacking parvalbumin+ somata but with processes extending towards the AC (Figures 5A-5C). Crucially, in 14-dpf fish, like adult *Danionella* (Figures 4B-D), Dc was at the dorsal surface, lateral to a calretinin+ area corresponding to Dm (Figure 5A). However, Dc was beneath Dl and Dm at caudal levels in 21-dpf, 28-dpf, and adult zebrafish (Figures 4A, 5B and 5C).

**Figure 5.**
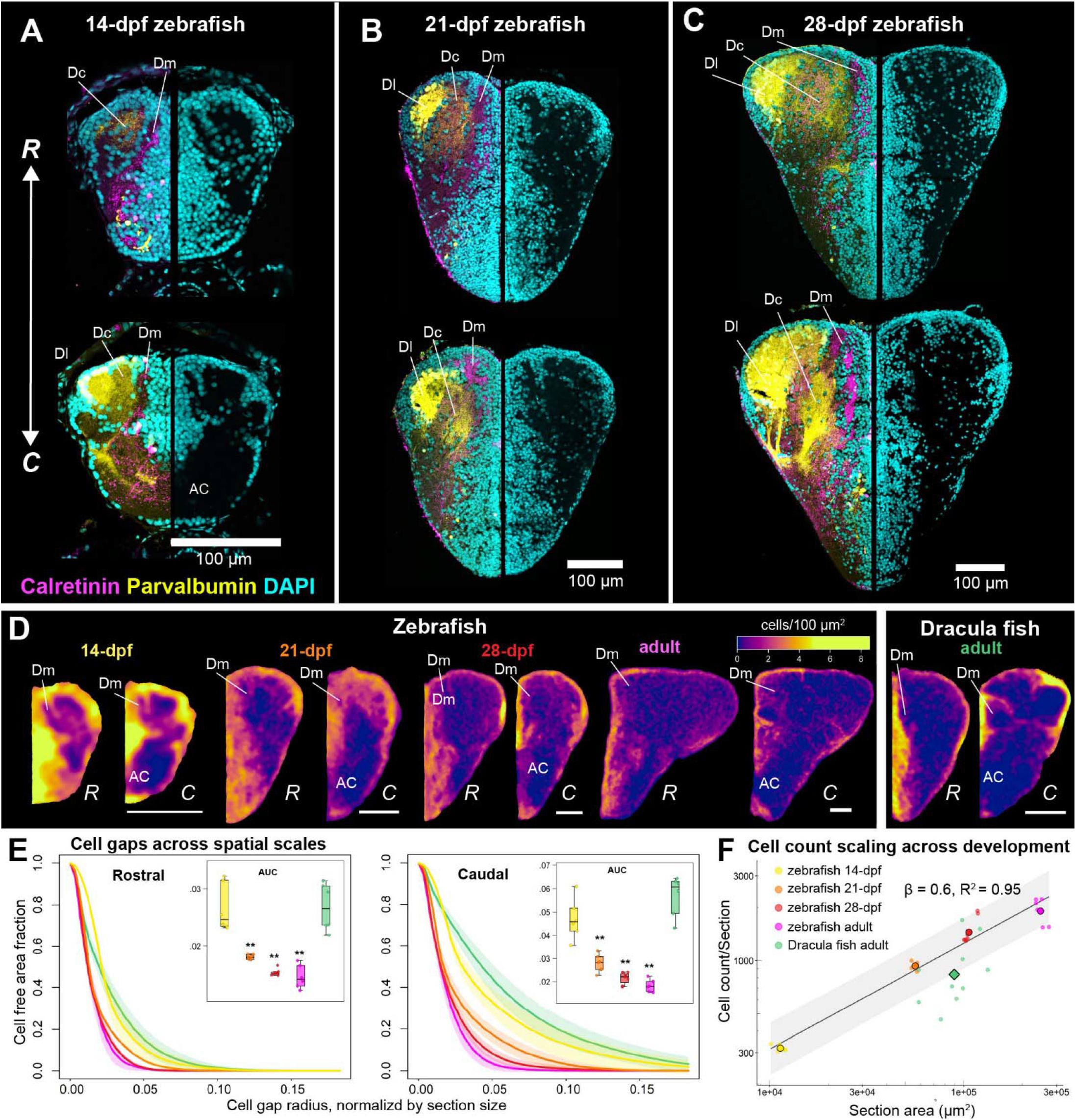
Comparison with zebrafish across development supports paedomorphic retention of larval telencephalic cytoarchitecture in adult Dracula fish. (A-C) Transverse sections show calretinin (magenta), parvalbumin (yellow) and DAPI (cyan) in telencephalon of (A) 14-dpf, (B) 21-dpf and (C) 28-dpf zebrafish at rostral (*R*) and caudal (*C*) levels. (D) Hemisection cell density heatmaps at rostral (*R*) and caudal (*C*) levels across developmental stages in zebrafish compared to adult Dracula fish (n = 6 per map). (E) Cell free area fraction across increasing gap radii, normalized to hemisection size (√ area) at rostral and caudal levels. Inset boxes show area under the curve (AUC) comparisons. Adult, 28-dpf and 21-dpf, but not 14-dpf zebrafish, differ significantly from adult Dracula fish (p < 0.01, Cliff’s δ = 1, Mann-Whitney U test). (F) Cell count per-hemisection over section area (log-log). Line of best fit characterizes zebrafish developmental progression. Larger color dots with black outlines reflect group means. Dracula fish adults (green diamond) have 0.71x fewer cells per-section (0.60-0.85% CI, p < 0.001) than equivalently sized zebrafish sections (between 21- and 28-dpf).

### *Danionella* adults and 14-dpf zebrafish larvae share compartment-like caudal pallium

Like *Danionella* adults (Figures 4B-4D), the caudal pallium of 14-dpf zebrafish appeared compartment-like (Figure 5A). Parvalbumin+ somata in Dl were restricted to the dorsal periventricular zone and cell bands corresponding to the apparent borders of Dm, Dl and Dp. Otherwise, the caudal pallium was largely devoid of cells. Parvalbumin+ and calretinin+ fibers filled the central neuropil of each caudal division, like *Danionella* adults. By 21-dpf, the caudal pallium was more diffusely filled with DAPI-labeled cells, and parvalbumin+ somata in Dl were displaced ventrally from the dorsal surface. At 28-dpf, these parvalbumin+ somata were more widely distributed throughout Dl. Pallial cytoarchitecture and parvalbumin and calretinin labeling in 28-dpf zebrafish resembled adult zebrafish (Figures 5A-5B, S4A, 4A and S3A).

To quantify these differences, we generated density heatmaps of segmented cells from rostral and caudal telencephalon hemisections across zebrafish developmental stages and compared these to zebrafish and Dracula fish adults (Figure 5D, n = 6 per level per group). This confirmed a population level developmental pattern in zebrafish with interior pallial zones largely devoid of cells at 14-dpf, then filled in at 21-dpf and older. We plotted the cell-free area fraction as a function of increasing radius, normalized to the hemisection area (Figure 5E). Curves that lie higher and extend to larger distances indicate larger cell-free regions. The zebrafish curves, ordered by age, showed 14-dpf fish higher with a more rightward shifted curve. The area under the curve for adult Dracula fish was significantly higher than adult, 28-dpf and 21-dpf, but not 14-dpf zebrafish (δ = 1, p < 0.01 for all significant differences, Mann-Whitney U tests with Holm correction). In contrast, Dracula fish telencephalon sections were closest in size to 28-dpf zebrafish (Figures S4B, S4C).

Examining the relationship between cell count and section size, zebrafish showed allometric scaling across developmental stages (β = 0.6, R^2^ = 0.95). Dracula fish adults fell significantly below this relationship, with 0.71x the cells at a given section area (0.60-0.85, 95% CI, p = 0.00044, ANCOVA) and thus had an average cell count per section closer to 21-dpf zebrafish (Figures 5F and S4D).

### Rostrocaudal partitioning of *Danionella* telencephalon

Calretinin and parvalbumin parsimoniously delineated all divisions of the Dracula fish telencephalon identified and further suggested a fundamental rostro-caudal partitioning of pallial subdivisions (Figures 6A and 6B, Video S1). These patterns were largely similar in *D. cerebrum*‘s telencephalon (Figure S5), suggesting a genus-wide phenotype.

**Figure 6.**
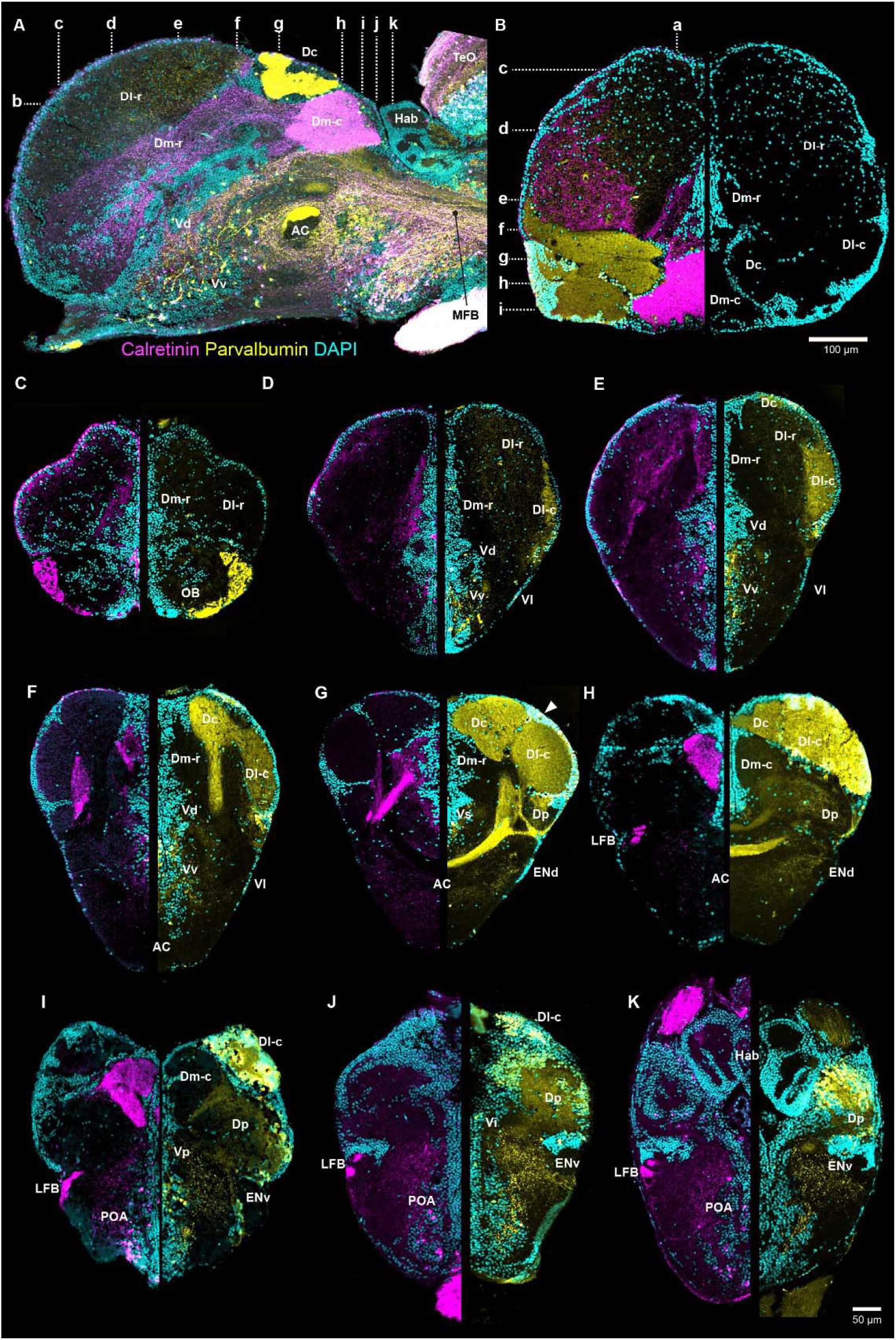
Rostrocaudal partitioning of Dracula fish telencephalon. (A-K) Sagittal (A) horizontal (B) and transverse (C-K) sections showing calretinin (magenta), parvalbumin (yellow) and DAPI (cyan) across Dracula fish telencephalon. Dl-c rostral to AC (D-F) is continuous with Dl-c at AC (G,H); however, in horizontal sections, a band of parvalbumin+ cells separated rostral Dl-c from caudal Dl-c (see Figure S6B). At caudal levels of Dl-c, a dorsal wedge of cells appeared at the border with Dc (arrowhead in G) which expanded caudally into a subcompartment-like structure before becoming indistinguishable from Dl-c (H). See also Figure S6C. Dc does not appear until midway through the telencephalon (E, F). In zebrafish adults, the dorsal portion of Dc, as defined by parvalbumin+ fibers, continues further rostrally, almost to the olfactory bulbs (Figure S3A, also^104^). Lowercase letters and dotted lines in (A,B) indicate approximate position of consecutive transverse sections (C-K).

#### OB and pallium

OB glomeruli were among the densest telencephalic regions labeled with both markers; labeling elsewhere in the OB was negligible (Figure 6C). Calretinin+ fibers were especially dense caudal to the AC, allowing us to partition a calretinin-rich caudal Dm subdivision (Dm-c) from a rostral subdivision (Figures 6A and 6B). Calretinin label first appeared at the dorsolateral periventricular surface (Figure 6C) and then expanded centrally (Figures 6D and 6E), before forming a dense lateral band (Figure 6F). This band and one emerging from Dm-r fed into the lateral forebrain bundle (LFB; Figures 6H-K), which was continuous with the diencephalon’s preglomerular complex (Figure S6A, Video S1).

Dense parvalbumin+ fibers and cells distinguished a parvalbumin-rich caudal (Dl-c) subdivision from a rostral (Dl-r) one essentially lacking parvalbumin (Figure 6B). Dc first appeared as a thin parvalbumin+ fiber field in the dorsal periventricular zone at mid-rostral levels (Figure 6E), before forming an expansive pallial division (Figures 6F and 6G). Both Dc and Dl appeared to contribute to the thick bundle of parvalbumin+ fibers crossing the AC (Figures 6G, 6H and S3B). The highest density of parvalbumin+ cells was in the caudal poles of Dl-c and Dp (Figures 6I-6K); Golgi-like labeling showed Dl neuronal processes projecting into its neuropil (Figure S6D).

#### Subpallium

Within V, parvalbumin+ cells were more common than calretinin+ ones and, mirroring *islet1* distribution (Figures 2I-2L), extended from Vv to Vs (Figures 6D-6G). Calretinin+ cells were distributed throughout Vv, while a separate population tightly clustered along the midline tracked the border between Vd and Vv, and into Vs (arc of cells in Figure 2A, Video S1). Calretinin and parvalbumin labeled cells were scattered in Vp. Vl and ENd contained calretinin+ cells, while ENv cells were parvalbumin+ (Figure S6Ei-iii).

### 3D atlas of Dracula fish telencephalon

To serve as a resource for functional imaging, we created a 3D atlas of the Dracula fish telencephalon and generated a transgenic Dracula fish line expressing nuclear-localized GCaMP6s under the *elavl3* promoter. Consistent with expression patterns in *D. cerebrum*^34,39^, GCaMP appeard widespread throughout the adult brain (Figure 7A). The telencephalons from nine fixed, cleared brains were imaged at high resolution (∼0.5 µm lateral, 2 µm z-steps) with an Airyscan confocal system (Figure 7B) and averaged into a reference volume using the ANTs iterative registration package (Figure 7C). Based upon the cumulative evidence from molecular markers, olfactory tracings and species comparisons (Figures 2-6), we annotated 15 telencephalic regions onto the reference volume (Figure 7D, Table S1). To demonstrate the utility of this atlas, we performed 2-photon imaging of spontaneous calcium activity across the telencephalon of a male Dracula fish, sequentially imaging 32 planes at 10 µm intervals, covering ∼85% of its depth (Figures 7E and 7F). We registered this individual functional imaging volume to the segmented atlas, extracted calcium traces with Suite2p from 6355 neurons and assigned them to brain regions (Figures 7G and 7H). We calculated mean ΔF/F and transient rate per brain region (Figures 7I and 7J). We identified neurons in the top 10% across the entire dataset, then plotted the fraction of cells for each region that were in this top 10% (Figure 7K). Dc was the most enriched for high mean ΔF/F (0.235) while OB (0.299) and Dl-r (0.252) were particularly enriched for high transient rates.

**Figure 7.**
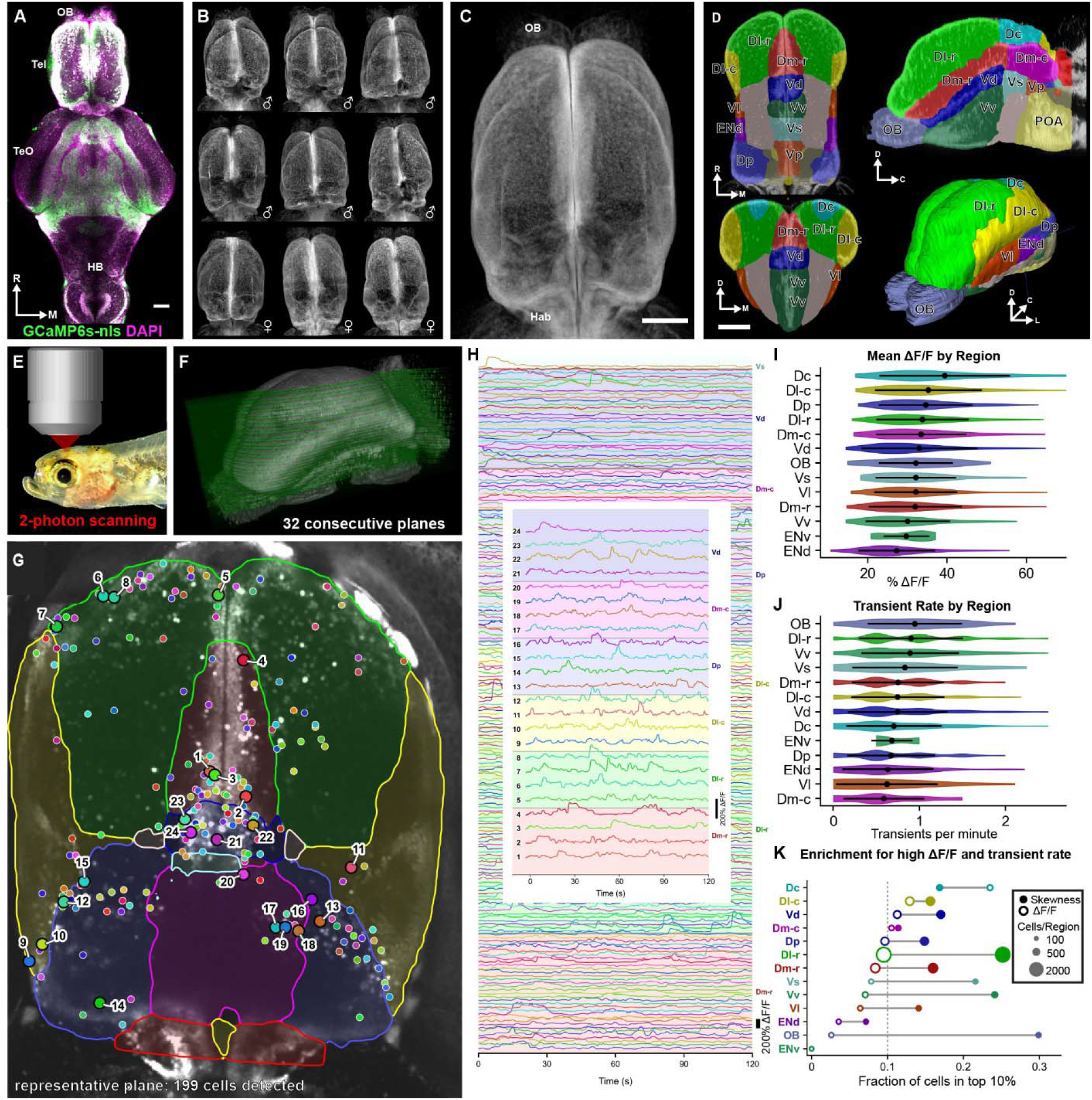
Development of 3D telencephalon atlas and pipeline for registration of functional imaging. (A) Transgenic adult Dracula fish brain expressing pan-neuronal nuclear-localized (nls) GCaMP6s (green), labeled with DAPI (magenta). (B) Telencephalons from nine GCaMP individuals. Intensity projections from fixed, cleared brains. (C) Averaged reference brain created from volumes shown in (B) using ANTs. (D) Segmented brain regions shown in horizontal, sagittal, coronal and 3D views. (E) Paradigm for proof-of-concept alignment of functional imaging data. Awake, immobilized GCaMP Dracula fish male under 2-photon excitation. (F) Spontaneous calcium signals recorded sequentially across 32 planes (10-micron steps). (G) Representative single plane with reference brain region boundaries warped onto individual brain. Colored dots indicate segmented GCaMP nuclei from spontaneously active neurons. (H) Calcium traces from 199 segmented cells in (G). Background colors and right axis labels indicate brain regions. Inset: 24 representative activity traces with numbers indicating location of cells in (G). (I) Mean ΔF/F of neurons per region, recorded across all 32 imaging planes (n = 6355). (J) Mean transient rate of neurons, per region. In (I) and (J) black circles are means and lines are standard deviation. (K) Proportion of cells, by region, within the top 10% of global values for mean ΔF/F and transient rate. Vertical dotted line indicates chance level (10%). Size of circles reflects number cells in that region.

## DISCUSSION

The size of nearly all adult vertebrate brains limits the scale at which brain-wide networks can be studied during ongoing behavior. *Danionella* fishes are currently the only adult vertebrates where size, optical transparency and genetics allow for multi-scale observation and perturbation of neuronal circuits throughout the entire brain.^23,26,29,30,36,39^ This advantage is afforded by progenetic paedomorphosis, a form of heterochrony resulting in miniature, sexually mature adults that retain larval or juvenile traits.^48,49,109^ Focusing on Dracula fish, we report a reorganization of brain regions that are proposed homologs or analogs of mammalian neocortex (Dc) and pallial amygdala (Dm).^47,66–68,103,110^ The larval-like telencephalon of Dracula fish adults, corroborated in other *Danionella* species, took us by surprise given the complexity of their adult behaviors. For example, as shown for Dracula fish^25^ and *D. cerebrum*,^23^ aggressive encounters include prominent acoustic displays by males (females are non-sonic) and, unique to male Dracula fish, a dramatic unfurling of a hypertrophied, fanged lower jaw^17,18,25,51^. Zebrafish do not begin to fight and court until close to the juvenile-adulthood transition^111–114^, well past the 14-dpf stage that most closely resembles the adult *Danionella* telencephalon.

Consistent with *Danionell*a’s organism-wide paedomorphic phenotype^15–18,50,51^, our results show that evolutionary shifts in developmental timing can lead to novel patterns of adult vertebrate brain organization, which in this case challenge the widely held link between elaboration of the adult brain and behavioral complexity in vertebrates^115–118^.

Due to a dramatic reduction in cell migration from ventricular zones in adult *Danionella*, we could not readily transpose the cytoarchitectonic details recognized for nearly the last 65 years in comparative studies of teleosts ^62,94,119^, including detailed brain atlases available for zebrafish.^38,55^ Fortunately, we could show by using conserved molecular markers broadly indicative of pallial and subpallial regions across vertebrates and of specific divisions in zebrafish and other teleosts (Table S1) that although the adult *Danionella* telencephalon has a larval-like cytoarchitecture resembling 14-dpf zebrafish, it retains most telencephalic regions common to adult teleosts, including zebrafish. Our comparative framework, annotated 3-dimensional atlas, and transgenic GCaMP Dracula fish line make possible the direct testing of the neural circuit mechanisms of differences in social behavior between *Danionella* species. Furthermore, these resources can aid in explaining how evolution and development of telencephalic complexity, or lack thereof, regulates seemingly complex social and other behaviors manifested in adulthood.

### Paedomorphic phenotype of *Danionella* telencephalon

*Cell migration*. During vertebrate brain development, most neurons are born from progenitors originating in periventricular zones, then differentiate and move via active migration or passive displacement to their mature locations.^120^ The reduced cellular density distal from the periventricular zones in *Danionella* telencephalon, particularly caudal pallium, suggests these processes are arrested. Strikingly, its caudal pallium resembles that of 14-dpf zebrafish, namely compartmental-like divisions of dense neuropil delimited by dense bands of cells (see Figures 1G, 4B-D, 5A, 5D). We show in zebrafish that a prominent migration of cells into the caudal pallial neuropil occurs between 14 and 21-dpf. The specific developmental processes arrested in *Danionella* could include neurogenesis rates, migration of neurons along radial glia, or passive tissue changes affecting morphogenetic movements, such as composition of extracellular matrix.^121^

#### Recognizing subpallium and pallium

The results for Dracula fish are consistent with the impression that organization of the teleost subpallium is more evolutionarily conserved than that of the pallium.^1,45,47,62,122–125^ *Dlx2a*, *islet1*, *otpa*, NPY, TH and calretinin expression in adult Dracula fish identified the pallial (D)-subpallial (V) boundary and midline-positioned subpallial nuclei (Vd, Vv, Vs, Vp, Vi). Topography and cytoarchitecture helped identify three of the four migrated subpallial populations (Vl, ENd, ENv). NPY helps distinguish Vl from ENd. A recent zebrafish study reported Vl as NPY-negative;^71^ however, we found NPY+ cells in Vl of both Dracula fish and zebrafish (Figures 2M and S2E). Previous work^78,126–128^ shows NPY+ cells extending from Vl into ENd (often designated EN). Unlike zebrafish,^71,78^ we did not observe NPY+ cells in Dracula fish Vd. This is unlikely due to paedomorphosis, as the Vd of 6-dpf zebrafish has NPY+ cells.^129^

The only subpallial region we did not identify was Vc. In contrast to other migrated subpallial regions discernable with molecular markers in 3-dpf zebrafish (Vl, ENd, ENv),^5^ Vc is reported only in adult zebrafish^55,93^ and other teleosts.^62,76,94,130^ In closely related goldfish (*Carassius auratus*), some recognize a distinct Vc,^76^ while others consider it continuous with Vd.^95^ Zebrafish Vc is hypothesized to migrate from larval Vd.^87,93^ We note a candidate Vc appearing first in 28-dpf zebrafish as a cell cluster slightly separate from Vd (Figure S4E). No molecular markers are known to distinguish Vc from Vd.^71,86,87,93,131^ A recent study in adult zebrafish using transgenic driver lines identified a population of *penkb+* neurons in Vc projecting to Vl, but Vc and Vd were grouped together as a proposed homologue of the mammalian striatum.^71^ A Vc might be identifiable in Dracula based on *penkb* expression and connectivity to Vl, despite its apparent absence due to reduced cell migration wrought by paedomorphosis.

Calretinin and parvalbumin labeling demarcated rostral and caudal divisions of Dm and Dl relative to the AC. Comparable variation has long been recognized^45,62,130^ and underscored by recent connectivity and single-cell transcriptomic studies.^69,70,76,108^ The only major pallial divisions we did not find in Dracula fish were Dd and nucleus taenia (NT). Dd is recognized as a division of densely packed cells between Dm and Dl^45,62,130^, and NT as a dense band of cells adjacent to Dp.^45,62,94,130^ Zebrafish Dd appears indistinguishable from Dl using cytoarchitecture, connectivity and immunohistochemical markers.^69,78^ Consequently recent studies consider it a subdivision of Dl^69^ or do not recognize it.^38,104,108^ Like Dd, no identified molecular markers distinguish NT from Dp, and many contemporary zebrafish studies do not consider it.

#### Topographic reorganization of pallium

Dc, originally defined as a deep, large-celled region in ray-finned fishes,^45^ resides at the dorsal surface in adult Dracula fish. Its correct identification is crucial as it has been proposed as homologous in whole ^68,104^ or in part^61,132^ to the dorsal pallium of tetrapods, which in mammals includes the neocortex^1^. Alternatively, some consider Dc a subdivision of one or more other pallial divisions.^47,67,76^ In adult zebrafish, Dc exhibits dense parvalbumin+ fibers but not neurons,^68,103^ co-expression of *eomesa* and *emx3,*^107^ and little-to-no calretinin, GAD, NPY, or TH label.^69,78^ We demonstrate all of this in Dracula fish Dc. Although a prior study in adult zebrafish reports *emx2* labels caudal Dc,^107^ we found *emx2* in Dp in Dracula fish. The pallial *emx2*+ region previously recognized in adult zebrafish as Dc is likely Dp (compare Figure 2C in^107^ showing *emx2* with Figure 18C in^69^ showing OB backfill of Dp).

Key to our study was a comparative approach using other *Danionella* species and 14 to 28-dpf larval zebrafish. Based upon parvalbumin and calretinin, we show Dc in a dorsal position in *D. cerebrum*, *D. priapus*, and 14-dpf larval zebrafish. It reaches its adult position in zebrafish beneath Dl and Dm by 21-dpf (Figures 4 and 5). A recent *D. cerebrum* brain atlas identifies a much broader rostral Dc and a caudal Dc in a deeper, adult zebrafish-like position based solely on expression of *eomesa*^133^, whereas we provide multiple lines of evidence to confirm Dc’s location and extent. *Danionella*’s Dc therefore retains a larval-like position, i.e., is paedomorphic. This resonates with the hypothesis that Dc in zebrafish arises from its own ventricular region, with the caudal end overgrown during development by Dl and Dm, while the rostral region remains at the dorsal surface.^104^ A subsequent study showed some Dc neurons clonally linked to neurogenic radial glia in the ventricular layer of adult Dm, while another subset of Dc neurons was not associated with clones in any part of the adult ventricular layer.^7^ It remains possible that in zebrafish, deep Dc is generated both from the ventricular layer of Dm and from an early ventricular layer of progenitors that become quiescent and covered during development. Our combined results from *Danionella* and zebrafish support this interpretation, but this should be confirmed with lineage tracing.

In Dracula fish, Dm sits beneath anterior Dl, while caudally it is deep to Dc. By contrast, Dm lies at the dorsal surface in other teleosts. We identified Dm based on calretinin+ fibers, a hallmark across teleosts,^76–78,134–136^ and high *emx3* but no *eomesa* expression.^103,107,137^ Interestingly, Dm in adult *D. priapus* and *D. cerebrum* has a position comparable to other teleosts. Across the zebrafish developmental stages we examined, Dm maintains a position similar to adults so its location in Dracula fish is unlikely directly linked to paedomorphosis. Miniaturization and associated heterochronic processes are known to constrain cranial morphology.^109,138^ The hypertrophied lower jaw of Dracula fish males is unique among *Danionella* species.^17,18^ Although the position of Dm in Dracula fish did not differ by sex, its development may involve craniofacial rearrangements, preceding sexual differentiation, creating lateral compression of the telencephalon and the derived position of Dm.^133^ Alternatively, other craniofacial differences between *Danionella* species, such as relative size or position of the eyes or inner ear, could contribute to the unique position of Dm in Dracula fish.

### *Danionella* telencephalon is a mosaic of larval- and adult-like characters

Paedomorphy is described in *Danionella* as organism-wide, but is nevertheless a mosaic phenomenon.^17,18^ While developmental truncation explains the absence of many bones in Dracula fish, the Weberian apparatus, a set of bones enhancing high frequency hearing, develops much earlier than in zebrafish and other Otophysi.^50^ A recent whole-body single-cell transcriptomic study in *D. cerebrum* found a mixture of larval- and adult-like gene expression in epithelial and connective tissues.^139^ The *Danionella* telencephalon is likely also a larval-adult mosaic. While the degree of cell migration in the caudal pallium and position of Dc suggest a 14-dpf zebrafish larval stage, cell counts in Dracula fish telencephalon are closer to 21-dpf zebrafish, while its size appears more similar to 28-dpf zebrafish (Figures 5 and S4B-D). In zebrafish, a late-developing lateral region of the pallium expands significantly from a tiny sliver between 14-dpf and 3-months^6^ and corresponds to at least part of the parvalbumin+ caudal Dl of adults. In adult *Danionella*, the caudal Dl appears at least as well developed as in 28-dpf zebrafish in terms of its size relative to other pallial divisions (Figures 4B-4D, 5A–5C, S4A).

Connectivity may be a particularly insightful anatomical correlate of when the nervous system reaches an adult stage of maturation. OB connectivity with Dp and habenula is established by 5-dpf in zebrafish.^96^ Not surprisingly, we find this connectivity in adult Dracula fish. What about later developing connections? The predominant source of sensory inputs to the teleost pallium is from the preglomerular complex (PG), a thalamic-like collection of nuclei in the diencephalon.^140–144^ The lateral division (PGl) projects mainly to Dl and anterior/medial divisions (PGa/PGm) project mainly to Dm.^69,76,145^ A zebrafish enhancer-trap line exclusively labeling PGl revealed projections to Dl are not detectable until around 60-dpf and continue developing until 3-months, close to the onset of sexual maturation.^146^ Developmental data on PGa/PGm projections to Dm in zebrafish is lacking, but our calretinin labeling in Dracula fish shows a prominent coupling between these two regions (Figure S6A and Video S1), suggesting the corresponding PGl to Dl projection is similarly well developed, but this requires further investigation. New, relatively high-throughput connectomic methods may provide an unbiased means to compare neural circuits in *Danionella* to those in zebrafish across development,^40,147,148^ while single-cell sequencing may yet reveal larval, adult-like or, more likely, a mosaic of transcriptional profiles.^149^

### Functional consequences of paedomorphic telencephalon: trade-offs and innovations

Reduced migration and simplification of the nervous system has been described in miniaturized, paedomorphic amphibians.^150–153^ Simplified telencephala are also found in many non-miniaturized animals such as bichirs,^62^ lungfish,^154^ caecilians^151^ and amphibians more generally.^152^ Studies of neurophysiological and behavioral consequences of this simplification are rare and largely limited to miniaturized plethodontid salamanders.^109,152^ In terms of brain size, number of neurons, degree of cytoarchitectural differentiation and neuronal morphology, plethodontids appear to have one of the simplest adult vertebrate nervous systems.^152^ Despite this, visual acuity and prey capture rival larger, non-miniaturized congeners. This likely reflects a trade-off prioritizing neural investment to the center visual field at the expense of peripheral vision. It also involves several innovations including a massive increase in the number of ipsilateral retinotectal projections, hypothesized to improve depth perception.^138,152^ Does the apparent simplification of the *Danionella* telencephalon portend comparable trade-offs or innovations?

The striking compartment-like appearance of the caudal pallium in *Danionella* fishes is evocative of insect glomeruli which exhibit a topographic separation of neuronal somata from a dense neuropil,^155^ hypothesized to minimize distance between axons and dendrites, thus saving space and reducing energy consumption.^156^ The neuropil-dominated divisions of the *Danionella* caudal pallium might similarly serve to optimize space for sensorimotor integration within a miniaturized head. This character could be seen in this light not only as paedomorphic retention of a larval-like phenotype, but as an evolutionary innovation with adaptive functions, akin to the Dracula fish hypertrophied lower jaw and tooth-like fangs, the evolution of which Britz and Conway proposed was facilitated through progenesis by an escape from developmental constraints.^17,50,51^ Connectomics may prove to be quite insightful and reveal, for example, cryptic circuit motifs within the pallium’s neuropil compartments^157^ that support computations orchestrating adult behaviors within the confines of a larval-like pallial cytoarchitecture.

### Proposed homologies with tetrapods

Despite diverging more than 350 million years ago, the similarity of subpallial organization between fishes and tetrapods appears remarkably conserved.^1,88,89^ Most lines of evidence support proposed homologues of subpallial regions in ray-finned fishes, which include teleosts like *Danionella* and zebrafish, to tetrapods as follows: Vd and Vc as striatal, Vl and ENd as pallidal components of the basal ganglia, and Vv as septum,^71,91^ with recent work proposing Vi as a homologue of medial amygdala.^101^ However, telencephalic eversion has bedeviled attempts to identify pallial homologues.^1^ Widely held comparisons between ray-finned fishes and tetrapods are Dl to hippocampus, Dm to pallial amygdala, Dp to piriform cortex, and Dc to neocortex.^66,68,103^ Significantly, recent alternative hypotheses hold one or more of these proposed homologies are instead instances of homoplasy.^110,158^ The ease of optical and genetic access in *Danionella*, and amenability for longitudinal imaging across development,^27,28,33^ combined with multi-omic spatial data, may provide much needed developmental and functional evidence to resolve debates of homology versus homoplasy between fishes and tetrapods.

### Limitations of the study

In this study, we assume zebrafish as a good representative of the ancestral Cyprinidae condition. Examining developmental data in outgroups would confirm this, especially within the superorder Ostariophysi, which includes *Danionella* and zebrafish. Including other paedomorphic cyprinids could help untangle the phenotypic contributions of developmental truncation from developmental novelty.^159–163^ To identify common brain regions, we selected molecular markers based on available data in zebrafish and other teleosts, but future species comparisons using spatial transcriptomics could provide an unbiased alternative approach. Although the size of *Danionella* makes combining labeling, whole brain clearing and imaging tractable, we opted to rely principally upon thin histological sections because: (1) this facilitated comparison with the vast prior literature on teleost neuroanatomy, (2) it afforded superior resolution to resolve boundaries between very small brain regions, (3) many antibodies, which provide essential information on innervation patterns, did not work in whole mounts, and (4) the expression level of several transcription factors required thin sections and high magnification for detection. Annotations of the 3D atlas were therefore done manually by matching cytoarchitectural patterns between thin sections from individual brains and the averaged reference volume. Future optimization of tissue clearing and expansion microscopy with select antibody markers, or generating transgenic reporter lines (e.g., for parvalbumin or calretinin), could provide population-level 3D innervation patterns to complement and extend the present work. We did not quantitatively compare male and female Dracula fish; however, such analyses may uncover sex differences, as reported for the telencephalon and other brain regions in *D. cerebrum.*^133^

## ACKNOWLEDGEMENTS

We thank Brian Miller, Eric Schuppe and Maximilian Bothe for assistance with PCR, Anson Chen and Ke Wong for help with brain dissections. We thank Brian Miller, Sophia Ranalli and George Tsimis for help with transgenics and egg injections. We thank Mykola Kadobianskyi, James Jaggard and Javier How for discussions on brain clearing and registration methods. We thank John McCormick and the Feschotte lab for providing zebrafish. We thank Melissa Warden for generous use of her confocal microscope, and Joseph Fetcho for use of his lab resources and supportive discussions. We also wish to thank Rose Tatarsky, Najva Akbari, Sarah Campbell and Midge Marchaterre for their past and continuing work to develop Dracula fish resources. Imaging data was acquired through the Cornell Institute of Biotechnology’s Imaging Facility (RRID:SCR_021741), with NIH funding for the shared Zeiss LSM 710 (1S10RR025502) and Zeiss LSM980 (S10OD036283) confocal microscopes. This work was supported by the NIH (NS128891) and NSF (IOS 1656664, 2443461).

## AUTHOR CONTRIBUTIONS

A.H.B. and J.T.P. conceived of the project. J.T.P. carried out the experiments. A.H.B. supervised the project.

A.H.B. and J.T.P. wrote the manuscript.

---

**Video S1. Calretinin immunolabeling in cleared Dracula fish forebrain (related to Figure 6)**

Confocal z-stack through rostral Dracula fish forebrain cleared and immunolabeled with calretinin (magenta). Arrowheads follow calretinin+ processes from fields in Dl-r, Dm-r and Dm-c through the lateral forebrain bundle (LFB) to the anterior and medial preglomerular nuclei (PGa, PGm) in the basal diencephalon (see Figure S6A). Yellow asterisk follows calretinin+ midline ventricular cells that demarcate the pallial (area dorsalis, D)/ subpallial (area ventralis, V) border (see Figures 2A, 2B and S2A). Labels Vd and Vv are validated at their specific positions; note that the Vd/Vv and D/V boundaries follow an ascending rostrocaudal arc (see Figures 2A and 6A).

## METHODS

### Key resources table

| REAGENT or RESOURCE | SOURCE | IDENTIFIER |
| --- | --- | --- |
| Antibodies |  |  |
| Alexa Fluor 488, donkey anti-mouse | ThermoFisher Scientific | A-21202 |
| Alexa Fluor 488, donkey anti-rabbit | ThermoFisher Scientific | A-21206 |
| Alexa Fluor 594, donkey anti-mouse | ThermoFisher Scientific | A-21203 |
| Alexa Fluor 594, donkey anti-rabbit | ThermoFisher Scientific | A-21207 |
| Alexa Fluor 647, donkey anti-mouse | ThermoFisher Scientific | A-31571 |
| Alexa Fluor 647, donkey anti-rabbit | ThermoFisher Scientific | A-31573 |
| Calretinin (rabbit, polyclonal, 1:500) | Swant | CR7697 |
| GABA (mouse, monoclonal, 1:500) | Swant | 3A12 |
| GAD 65/67 (rabbit, polyclonal, 1:500) | Abcam | ab11070 |
| Hu (mouse, monoclonal, 1:1000) | ThermoFisher Scientific | 16A11 |
| Islet1 (mouse, monoclonal, 1:100) | DSHB | 40.2D6 |
| Neuropeptide Y (rabbit, polyclonal, 1:500) | Millipore | N9528 |
| Parvalbumin (mouse, monoclonal, 1:500) | Millipore | MAB1572 |
| SV2 (mouse, monoclonal, 1:50) | DSHB | SV2 |
| Tyrosine hydroxylase (mouse, monoclonal, 1:500) | Millipore | MAB318 |
| Biological samples |  |  |
| cDNA from brain - *Danionella dracula* | This paper | N/A |
| Chemicals, peptides, and recombinant proteins |  |  |
| Neurobiotin | Vector Laboratories | SP-1120-20 |
| ProLong Gold Mountant with DAPI | ThermoFisher Scientific | P36931 |
| Streptavidin, Alexa Fluor 488 Conjugate | ThermoFisher Scientific | S32354 |
| EasyIndex RI 1.465 | Life Canvas Technologies | EI-500-1.46 |
| Critical commercial assays |  |  |
| HCR probes, buffers and amplifiers | Molecular Instruments | https://www.molecularinstruments.com |
| Experimental models: Organisms/strains |  |  |
| *Danio rerio* wildtype | Feschotte Lab, Cornell University | N/A |
| *Danionella cerebrum* wildtype | Bass Lab, Cornell University | N/A |
| *Danionella dracula* wildtype | Bass Lab, Cornell University | N/A |
| *Danionella dracula* Tg(elavI3-H2B:GCaMP6s) | Bass Lab, Cornell University | N/A |
| *Danionella priapus* wildtype | Bass Lab, Cornell University | N/A |
| Recombinant DNA |  |  |
| Tol2-elavI3-H2B-GCaMP6s | Freeman et al.<sup>166</sup> | Addgene plasmid #59530 |
| Software and algorithms |  |  |
| ImageJ/FIJI | Schneider et al.<sup>167</sup> | https://imagej.nih.gov/ij/ |
| UGENE | Okonechnikov et al.<sup>168</sup> | https://ugene.net/ |
| ANTs, ANTsPy | Tustison et al.<sup>169</sup> | https://github.com/antsx/antspy |
| Suite2p | Stringer et al.<sup>170</sup> | https://github.com/MouseLand/suite2p |
| Cellpose | Stringer et al.<sup>171</sup> | https://github.com/mouseland/cellpose |
| R v4.6.1, spatstat v3.6-1 | Baddeley et al.<sup>172</sup> | https://spatstat.org/ |

## EXPERIMENTAL MODEL AND SUBJECT PARTICIPANT DETAILS

### Danionella fishes

Adult *Danionella dracula* (Dracula fish) and *D. cerebrum* were maintained in breeding colonies at Cornell University following previously described husbandry methods.^23,25^ Since we collect eggs daily from breeding tanks, we could ascertain the age of these two species. All Dracula fish and *D. cerebrum* used in this study were between 8 months and 1.5 years old, which is well past the previously reported onset of sexual maturity.^23,25,31^ We were unable to breed *D. priapus* in the lab. Wild-caught specimens were purchased from a commercial supplier (The Wet Spot Tropical Fish, Portland, OR) and maintained under similar conditions as Dracula fish. Although we could not determine the age of *D. priapus*, they were examined to ensure maturity. All three *Danionella* species were sexed by visualizing gonads through the transparent abdominal cavity under a dissecting microscope and noting the position of the urogenital pore. Females have mature oocytes and a urogenital pore anterior to the anal fin. Males have testes and a urogenital pore between the pelvic fins. We report the average standard length (for adult fishes, snout to caudal peduncle of the tail) and sex for all subjects in Table S2.

### Staging of zebrafish

Wildtype *Danio rerio* (zebrafish) eggs were collected from breeding tanks and raised in age-matched cohorts in a zebrafish facility at Cornell University. Although days-post-fertilization (dpf) is the predominant metric reported across most zebrafish neurobiological studies, body size is a far better correlate of developmental milestones than chronological age^165^ because developmental rate is influenced by environmental and genetic (i.e., strain) factors. We therefore report standard length (as defined across stages in^165^) for all zebrafish age cohorts used in this study (Table S2). We adopt definitions for zebrafish stages which broadly agree that the juvenile stage begins following the development of most adult characters (scales, full complement of adult fin-rays), but in the absence of sexual maturity (∼30-45 dpf), while the adult stage begins with the onset of sexually mature gametes and reproductive behaviors (∼90 dpf).^14,165,173–175^ Based upon these criteria, the oldest pre-adult zebrafish age cohort we examined (28-dpf) would still be considered larval or late-larval, on the cusp of reaching the juvenile stage.

## METHOD DETAILS

### Brain tissue preparation and sectioning

All subjects were deeply anaesthetized in 0.025% benzocaine dissolved in aquarium water, followed by assessment of standard length and sex (where determinable, see above). Brains were exposed by removing the skin (and bone for older zebrafish) over the dorsal surface and then postfixed overnight at 4° C with gentle agitation. A subset of Dracula fish (N=25) were transcardially perfused with ice-cold freshwater teleost Ringers solution followed by 4% paraformaldehyde (PFA) in chilled 0.1 M phosphate buffered saline (PBS), followed by overnight fixation in 4% PFA. For all samples, except 14-dpf zebrafish larvae, brains were washed the next morning 3x 20 min in PBS, dissected out into PBS, and then cryoprotected by immersion in a graded sucrose solution series (15%, 30% w/v in PBS). Zebrafish larvae at the 14-dpf stage were treated similarly, except brains were not dissected out and sectioned *in situ*. Samples were immersed in biopsy molds filled with Tissue-Tek O.C.T. (Sakura Finetek USA, Inc.) and frozen in a slurry of crushed dry ice and 100% ethanol. Frozen, embedded samples were stored at −80° C until sectioning. All samples were sectioned at 12 µm on a cryostat (Cryocut 1800, Reichart-Jung) held at −20° C and collected onto Superfrost Plus slides (ThermoFisher Scientific). Most samples were collected on a single series, but a subset of *Danionella* and all adult zebrafish were collected in two alternate series. All slides were stored in sealed containers at −80° C until further processing. We found that while transcardial perfusion improved the odds that a *Danionella* brain retained near perfect tissue morphology after sectioning, immersion fixation produced excellent results as well, ∼80% of the time. When tissue morphology was good (as evaluated by the cytoarchitectural detail provided by DAPI), we saw no difference in signal quality of several antibodies (calretinin, parvalbumin, tyrosine hydroxylase) and *in situ* probes (*dlx2a*, *eomesa*). Two key factors for preserving *Danionella* brain tissue integrity for cryostat sectioning were minimizing time in sucrose solutions (samples were transferred as soon as they sunk to bottom) and rapid freezing with 100% ethanol and dry ice.

### Immunohistochemistry on sections

Samples on slides were allowed to thaw at room temperature (RT). A hydrophobic border was then drawn around the perimeter of the slide (ImmEdge Pen, Vector Laboratories). Slides were washed 2x 10 min in PBS, blocked for 1 h (10% normal horse serum, 0.3% Triton X-100 in PBS) and incubated with primary antibodies (in blocking solution, see STAR Methods for antibody dilutions) overnight. Slides were washed 6x 10 min in PBS + 0.5% horse serum and incubated with fluorescent secondary antibodies (1:200 dilution in blocking solution) for 2 hours. Slides were then washed 3x 10 min in PBS and coverslipped with ProLong Gold with DAPI (ThermoFisher Scientific). Only one antibody (*islet1*) required antigen retrieval, which was performed by incubating slides for 15 min in 150 mM Tris-HCl at pH 9.0 warmed to 70° C prior to blocking.^176^

### Whole mount clearing and immunohistochemistry

Dissected Dracula fish brains were cleared using DEEP-Clear.^177^ Brains were incubated overnight in prechilled acetone at −20° C, then cleared for 24 hours in solution 1.1 (pH = 11) at 37° C with constant agitation. Following PBS washes, brains were refractive index matched in EasyIndex (RI = 1.465, LifeCanvas Technologies). For imaging, brains were mounted in 1% low melt agarose dissolved in EasyIndex. For antibody labeling, prior to immersion in EasyIndex, brains were treated with antibodies (calretinin and anti-rabbit AlexaFluor 594) as described for sections on slides; however, incubations were extended to 7 days at 4° C with constant agitation. All steps were carried out in 1.5 ml tubes in 500 µL of solution.

### *In situ* hybridization on sections

To design probes, we first BLAST searched the orthologous genes from zebrafish against a scaffold level Dracula fish genome using UGENE^168^. We manually inspected the Dracula fish contigs to confirm exon/intron boundaries and designed primers to amplify 500-800 bp sections for each gene, including presumptive UTRs. We extracted mRNA from pooled Dracula fish brain tissue, synthesized cDNA and verified gene targets with Sanger sequencing. Hybridization chain reaction (HCR) probes, hairpin amplifiers and buffers were purchased from Molecular Instruments. HCR v3.0 probes and reagents were used for *dlx2a* and *otpa*, while HCR Gold probes and reagents were used for *eomesa*, *emx2*, and *emx3*. Care was taken during probe design to omit conserved sequences from all paralogs (e.g., *dlx2b*, *otpb*, *emx1*). Staining was carried out according to manufacturer’s published protocols for performing HCR on tissue sections on slides (“HCR RNA-FISH v3.0 protocol for fresh frozen or fixed tissue sections” and “HCR Gold RNA-FISH User Guide”).^178^

### Neurobiotin injections

Dracula fish (N=8) were anesthetized in 0.025% benzocaine dissolved in aquarium water and then secured with 2% agarose on a custom Sylgard petri dish filled with aquarium water. Neurobiotin (Vector Laboratories) was dissolved in 0.5 M KCl (5% w/v) and delivered via a pressure injector and micro-injection needles with a tip diameter of ∼10 µm. Neurobiotin was injected into the olfactory bulb unilaterally using a micromanipulator to advance the needle from the side at an angle approximately 60° from the rostral pole of the head, targeting a position between the nares and eye. After 1-2 injections (∼100 nL each), we removed the fish from agarose and allowed them to recover, under observation, in a petri dish with fresh aquarium water. Fish were then maintained in recovery tanks for 48 h, until collection of brain tissue (see Brain tissue preparation and sectioning). Slides with sectioned brain tissue were processed as described above (Immunohistochemistry), albeit with Streptavidin-Alexa Fluor 488 (1:500 diluted in blocking solution, ThermoFisher Scientific) to visualize the neurobiotin. To help identify pallial and subpallial brain regions, we co-labeled slides with calretinin and tyrosine hydroxylase, visualized with Alexa Fluor 647 and 594, respectively (1:200 diluted in blocking solution, ThermoFisher Scientific).

### Evaluation of anatomical markers and image acquisition

All stained slides were evaluated on a Zeiss Axioskop 2 Plus epifluorescence microscope to confirm consistent patterns of expression for each marker. At least two series for each marker were then imaged using confocal microscopy (Zeiss LSM 710, 800 or 980). Tiling and z-stacks were used as needed and for higher magnification (40x lens) images. For calretinin-labeled cleared, whole brains, the telencephalon was imaged with a 10x lens and voxel size of 0.62 x 0.62 x 3.88 µm. Composite images were generated in FIJI/ImageJ,^167^ using contrast/brightness adjustment, maximum projections of z-stacks, stitching and lookup table selection. Images were cropped and re-sized for figures using Photoshop C27.5 and Illustrator V30.3.

### Cell segmentation and spatial statistics

#### Imaging parameters

For each group (adult Dracula fish, adult zebrafish, 28, 21 and 14-dpf zebrafish) we selected 6 animals from the set of DAPI-labeled slides and imaged a rostral and a caudal section of the telencephalon. The rostral sections were taken at approximately half the distance between the caudal end of the olfactory bulbs and the rostral appearance of the anterior commissure. The caudal sections were taken at the anterior commissure with the rostral appearance of the preoptic area. All images were acquired with a 20x air objective (0.8 NA) at 0.6239 µm/pixel for all groups except 14-dpf zebrafish which were imaged at 0.3119 µm/pixel. All images were single acquisitions of a single optical plane at the center of the section except for adult zebrafish, which were imaged as tiles of 2-4 images, stitched together in Zen software and converted to tif files.

#### Cell segmentation, image registration and cell density heat maps

The right hemisphere of each section was traced and exterior pixels masked in FIJI/ImageJ. DAPI-labeled nuclei were segmented using the Cellpose-SAM model.^171,179^ and the centroids of all nuclei extracted in FIJI/ImageJ. Hemisections within each experimental group were registered to a single reference section using deformable registration (ANTsPy^169^, SyNRA transform, mutual information metric). The manually traced tissue outlines and segmented cell centroids were then mapped into the reference section’s coordinate space via the inverse deformation field. Hemisection cell density was computed by 2D Gaussian kernel density estimation (σ = 4 µm) on a common 512 × 512 pixel grid and averaged across sections within each group to produce raw average density maps (R, spatstat package).^172^ All further analyses (see below) were conducted using the unwarped, original hemisection outlines and cell centroids.

#### Relative cell density across hemisection depth

For each hemisphere, the distance of every cell centroid from the hemisection edge (midline and lateral walls) was computed (spatstat, *bdist.points* function). Depth was then expressed as the proportion of section area lying within that distance of the boundary, obtained by analytic erosion of the section polygon (spatstat, *erosion* function). This was evaluated at 200 equally spaced distances and interpolated to generate the curves in Figures 1J and 1K, so that profiles from sections differing several-fold in size are placed on a common axis running from 0 at the hemisection edge to 1 at the deepest interior point. Cell density across 20 bins (each 5% of the relative depth) was divided by the density expected if the same number of cells were distributed uniformly over the section, giving a relative density (fold-enrichment over uniform) that is unitless and independent of section area and cell count. For descriptive purposes, a linear slope was calculated for each curve and averaged per group (Figure S1G). A center-to-edge ratio was computed as the average relative cell density of the inner 7 bins over the outer 7 bins (both 35% each of section area, Figure S1H) and compared between adult Dracula fish and adult zebrafish (n = 6 animals per group, one hemisphere each) using two-sided Wilcoxon rank-sum tests with effect size reported as Cliff’s delta; rostral and caudal levels were analysed separately as pre-specified independent tests and p-values were not pooled across them.

#### Cell free area fraction

For each hemisection, a distance map was computed on a 512 x 512 pixel grid, and for every pixel contained within the section, the distance to the nearest cell centroid was determined (spatstat, *distmap* function).

Distances were normalized within each section by the square root of its area (r), so that the resulting curves are invariant to overall section size and comparable across groups differing several-fold. The cell-free area fraction V(r) was then defined as the proportion of tissue pixels lying farther than the scaled distance r from any cell. This is the complement to the empty-space function F^172^, evaluated on the size-normalized axis. Curves that remain high out to larger relative radii (r) indicate that more of the section lies far from any cell. The area under the curve (AUC) was used as a single scalar to compare Dracula fish to all other groups (n = 6 animals per group, one hemisphere each) using two-sided Mann–Whitney U tests with effect size reported as Cliff’s delta and Holm correction applied within section level.

#### Cell number-to-section size scaling

Cell centroids were counted and outline areas measured for each hemisection; counts and areas were log10-transformed per hemisection and averaged within animal across rostral and caudal levels. The scaling of cell number with section area was estimated by ordinary least-squares regression of log cell count on log section area across the zebrafish developmental series (14-dpf, 21-dpf, 28-dpf, adult). Dracula fish was compared to this relationship by analysis of covariance with a binary species term, fitting parallel lines and estimating the vertical offset of Dracula fish from the zebrafish scaling line as a fold-difference in cell number at matched section area.

### Generation of transgenic Dracula fish *Tg(elavl3-H2B:GCaMP6s*) line

Dracula fish court and breed most consistently in community tanks, spawning throughout the day. To collect eggs for transgenics, we established 9 breeding tanks (10-15 males, 20-30 females in static, non-recirculating 10-gallon tanks) and collected eggs by checking nests approximately every 45 minutes. Dracula fish egg clutches are lightly adhered together and so were gently separated with blunt forceps into a custom designed agarose mold. Single-cell stage eggs were injected with ∼100 nL of Tol2 mRNA (25 ng/ µL) and Tol2-elavl3-H2B-GCaMP6s plasmid (10 ng/ µL). The Tol2 plasmid was a gift from Misha Ahrens^166^. Larvae were screened to identify male founders and outcrossed with colony bred wild-type female fish in community tanks (1-2 males, 10-20 females). In F1 transgenic fish, and all subsequent generation, GCaMP nuclear labeling was strong and overlapped substantially with DAPI in dissected, cleared brains, indicating near pan-neuronal labeling.

### 3-Dimensional Telencephalon Atlas Construction

#### Image acquisition

Cleared brains from 9 *Tg(elavl3-H2B:GCaMP6s*) Dracula fish (6 males, 3 females) were embedded and coverslipped in glass-bottom wellplates and imaged on a Confocal Zeiss LSM 980 with a water-immersion 10x objective (0.45 NA, Olympus) in Airyscan mode. Telencephalon volumes were acquired by imaging the fixed GCaMP fluorescence at voxel size 0.4874 x 0.4874 x 2 µm. Zen Blue software (Zeiss) was used to stich tiles and apply Airyscan processing and joint deconvolution.

#### Template construction

Contrast was enhanced in FIJI with the CLAHE algorithm, volumes converted to NIfTI format, then passed through an initial pre-alignment stage with the ANTs ‘Similarity’ function. The average of the 9 volumes was used as the seed to generate a final reference volume in ANTs on a dual 64-core AMD Epyc 9554 server (3TB RAM, Red Barn) with the following call: *antsMultivariateTemplateConstruction2.sh -d 3 -n 0 -q 200x200x100x20 -f 12x8x4x1 -s 4x3x2x1 -c 2 -j 8 -i 4 -z avg_template.nii images_to_register.txt*

#### Template annotation

The reference telencephalon volume was manually segmented into anatomical regions using ITK-SNAP (v4.0.2), using the cytoarchitectural detail from pan-neuronal GCaMP nuclear labeling and informed by the immunohistochemical, HCR and neurobiotin labeling on thin coronal sections. Given this, and that the vast majority of teleost brain atlases are presented largely in the coronal plane, we annotated the telencephalon in this plane at every 10-20 sections of the volume. A Gaussian smoothing filter was applied in ITK-SNAP (2 x 2 x 2 voxels), followed by filling of small, unlabeled voxels using a nearest-neighbor algorithm, and a final anisotropic Gaussian smoothing (sigma = 24 voxels) along the anterior-posterior axis to correct for jitter introduced by the coronal tracing and interpolation.

### GCaMP functional imaging and registration to atlas with anatomical labels

#### Two-photon functional imaging

Adult male *Tg(elavl3-H2B:GCaMP6s*) Dracula fish were briefly and lightly anesthetized in 0.0025% benzocaine dissolved in fish system water. Fish were injected in the trunk with 200 nL of pancuronium bromide (0.4 µg/µL in teleost saline solution) using pulled glass needles and a pressure injector. They were then held in a 3D printed holding chamber (adapted from^33^), supported by 4% agarose around the trunk and perfused through the mouth and over the gills with a peristaltic pump (ESI MP2, Elemental Scientific) at 1 ml min^-1^ with oxygenated, temperature controlled (25° C) fish system water. Fish were given 15-20 minutes for benzocaine to wear off, then imaged with an Olympus FVMPE-RS multiphoton microscope. GCaMP was excited at 920nm with an Insight X3 laser (SpectraPhysics) under a 10x long working distance objective (0.6 NA, XLPLN10XSVMP, Olympus) with galvo scanning. Spontaneous calcium activity was recorded consecutively across 32 anatomical planes of the telencephalon, each separated by 10 µm z-steps, for the entire pallium and roughly 60% of the subpallium (512 x 384 pixel frame size, 1.4063 pixels/µm, 1.2287 Hz frame rate). At the end of each recording session, a higher resolution structural image was acquired at 0.3906 x 0.3906 x 5 µm voxels, spanning the entire telencephalon (100 z-planes, 500 µm range). For both functional and structural imaging, laser power was modulated as a function of depth.

#### Image processing and calcium signal extraction

Functional imaging files were converted from OIR to tiff, then processed in suite2p^170,180^ (v1.1.0) for motion correction and ROI detection. Frame rate, the decay time constant (τ = 3.8 s, nuclear-localized GCaMP6s^181^), and expected ROI diameter of 4 × 4 pixels were specified; all other parameters used suite2p 1.1.0 defaults.

Because the GCaMP indicator used was nuclear-localized, raw fluorescence (F) was used directly, without neuropil subtraction. For each ROI ΔF/F was computed as (F − F0)/F0 × 100%, with F0 defined as the rolling 8th-percentile baseline of F over a 40 s window. Traces were median-filtered with a 3-frame (∼2.4 s) kernel.

#### Registration of functional imaging planes to annotated atlas

The 32 functional imaging planes were registered to the structural stack from the same fish, which was in turn registered to the reference telencephalon volume. Functional-to-structural alignment (ANTsPy) proceeded in three steps: pairwise 2D rigid registration corrected residual stage drift between the independently acquired planes; a global 3D rigid-then-affine fit aligned the resulting functional volume to the structural stack; and a per-plane 2D translation correction (cross-correlation template matching) removed remaining depth-dependent distortion, reducing the mean residual from 21.1 px to 0.94 px across the 32 analyzed planes. Structural-to-template registration used 19 manually placed landmark pairs to fit a 12-degree-of-freedom affine transform, followed by deformable (SyN, cross-correlation metric) registration at 4 µm isotropic resolution, with registration quality assessed by deformation-field smoothness and visual inspection.

Region labels from the annotated template were mapped onto each functional plane’s own pixel grid by applying the inverse of the combined transform chain to each plane’s true 3D physical position; because the functional-to-structural transform includes a real ∼19° rotation between imaging axes, each native plane was treated as a thin, tilted 3D slab rather than a flat slice, and labels were resampled onto it with nearest-neighbor interpolation to preserve discrete label identity. This yielded a per-pixel region map for each of the 32 analyzed planes. Each suite2p identified ROI was assigned to whichever annotated region contained the largest number of its pixels (a plurality rather than strict majority, since a footprint spanning multiple regions need not have any single region exceed 50%). The fraction of the ROI’s pixels belonging to that region was recorded as a purity score: near 1 for ROIs lying entirely within one region and lower for those straddling a region boundary. ROIs whose plurality of pixels fell outside any annotated region were excluded.

#### Mean ΔF/F, transient detection and enrichment analysis

For each ROI, mean ΔF/F was computed as the average of its ΔF/F trace across the full recording. Significant transients were detected following the general approach of Dombeck et al. (2007)^182^. Events were required to exceed a multiple of the noise standard deviation (SD), with the false-positive rate empirically calibrated by applying the identical detection criteria to the sign-inverted trace, exploiting the fact that genuine calcium transients are strictly positive-going. Per-neuron noise (σ) was estimated as the difference between the trace’s median and its 15.87th percentile (=1 SD under a Gaussian approximation). Candidate transients were peaks exceeding 3σ in height and 1.5σ in prominence, separated by ≥4 frames (∼3.3 s, approximating one nls-GCaMP6s decay); this threshold was chosen by applying the identical detection procedure to the inverted trace — where any detected events are by construction false positives — and selecting the smallest multiplier yielding a dataset-wide false-positive rate ≤10% (K = 3σ, ∼4% observed). Transient rate is reported as significant events per minute, per cell.

To ask whether particular brain regions were disproportionately composed of highly active cells, a dataset-wide threshold was defined for each per-cell metric (mean ΔF/F and significant-transient rate) as its 90th percentile across all retained cells. For each region, enrichment was then the fraction of that region’s own cells exceeding this threshold; since 10% of all cells fall above it by construction, a region with no relationship to the metric is expected to show ∼10% of its cells in this top decile, and each region’s observed fraction was compared against that chance level to describe relative over- or under-representation of high responders.

#### Use of AI-assisted code editing

Portions of the Python and R code used for data analyses were drafted and edited with assistance from Claude Code (Anthropic). All code was reviewed, tested and validated by the authors.

---

**Figure S1.**
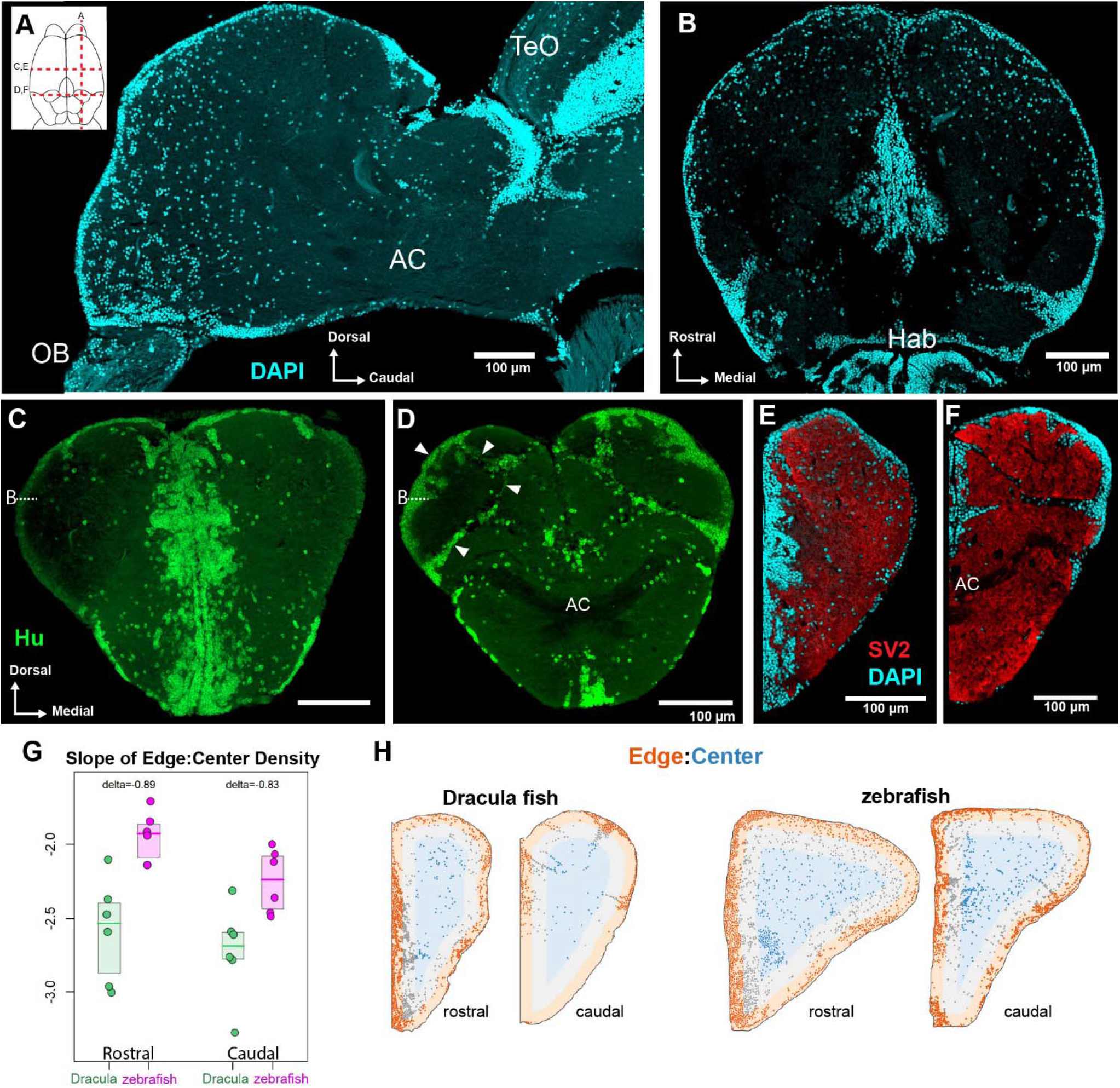
Cytoarchitectural characterization of the Dracula fish telencephalon (Related to Figure 1). (A) Sagittal and (B) horizontal sections through Dracula fish telencephalon showing cells labeled with DAPI (cyan). Note rostro-caudal gradient of decreasing cell density, highlighted in caudal pallium at level of compartment-like lobes (yellow arrowheads; also see Figures 1C, 1D). Top left schematic of dorsal telencephalon indicates approximate position of all sections. AC, anterior commissure; Hab, habenula; TeO, midbrain optic tectum. (C) Rostral and (D) caudal transverse sections showing neuronal somata labeled with Hu antibody (green). Cell-dense bands surrounding cell-sparse pockets (arrowheads, D) further highlight the lobe-like appearance of caudal pallium. B with dashed line indicates location of horizontal plane shown in (B). (E) Rostral and (F) caudal transverse sections showing synaptic vesicle protein 2 (SV) immunohistochemistry (red) and DAPI (cyan). (G) Comparison of relative cell density changes in adult Dracula fish (green) and zebrafish (magenta) telencephalon at rostral and caudal levels. Dracula fish have a steeper decline in cell density from the edge (midline and lateral periphery) to the center of each hemisphere, rostrally (Cliff’s delta = −0.89, p = 0.0087) and caudally (Cliff’s delta = −0.83, p = 0.0152). (H) Representative hemisection plots of cell centroids in Dracula fish and zebrafish telencephalon at rostral and caudal levels. Each plot from one individual. Corresponds to averaged heatmaps in Figure 1I. Analyses in Figures 1J and 1K compare the mean relative density of cell centroids in the center (inner third, blue) divided by the edge (outer third, red).

**Figure S2.**
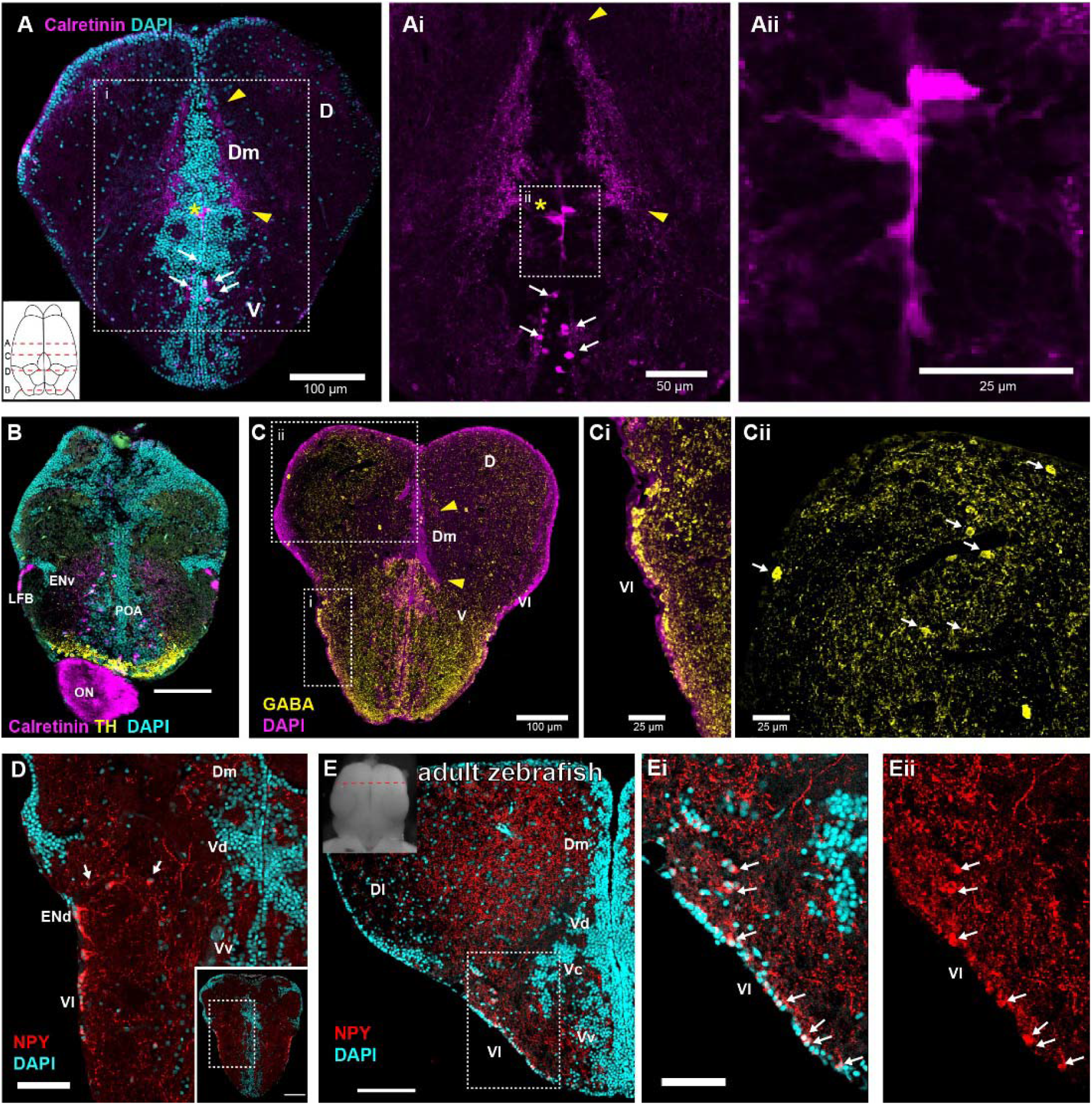
Immunohistochemical markers confirm divisions of D and V in Dracula fish telencephalon (Related to Figure 2). (A-D) Transverse sections; dorsal view schematic (A) indicates approximate levels. (A) Transverse section at comparable position to Figure 2B, showing calretinin immunohistochemistry (red) and DAPI (cyan). Yellow arrowheads mark calretinin+ fiber field of Dm, arrows mark calretinin+ cells in V, and yellow asterisk marks midline, ventricle-contacting calretinin cells. (Ai, Aii) Calretinin+ signal shown at increasing levels of magnification. Ventricular calretinin+ cells mark the precise boundary between pallium and subpallium, offering a potentially useful landmark when viewing the telencephalon in the horizontal plane (e.g., during calcium imaging). (B) Tyrosine hydroxylase (TH) antibody labeled cells (yellow) largely within far ventral preoptic area (POA). Calretinin antibody labeled cells (magenta) are distributed throughout preoptic area (POA). Note calretinin+ label within the lateral forebrain bundle (LFB) lateral to the ventral entopeduncular nucleus (ENv). Cell nuclei labeled with DAPI (cyan). (C) Transverse section showing GABA immunohistochemistry (yellow) and DAPI (magenta). Hatched boxes indicate highlighted regions of GABA+ cells in Vl (Ci) and D (arrows, Cii). (D) Neuropeptide Y (NPY) antibody labeled cells (red) just rostral to the anterior commissure in dorsal entopeduncular nucleus (ENd) and Vl. Note medial NPY cells (arrows) between ENd and Vd. Cell nuclei labeled with DAPI (cyan). (E) Transverse section from adult zebrafish rostral telencephalon shows NPY labeling (red). Inset dorsal view of telencephalon indicates approximate location of section. Small dotted white box denotes closeup of NPY cells (arrows) in Vl shown with DAPI (cyan, Ei) and without (Eii).

**Figure S3.**
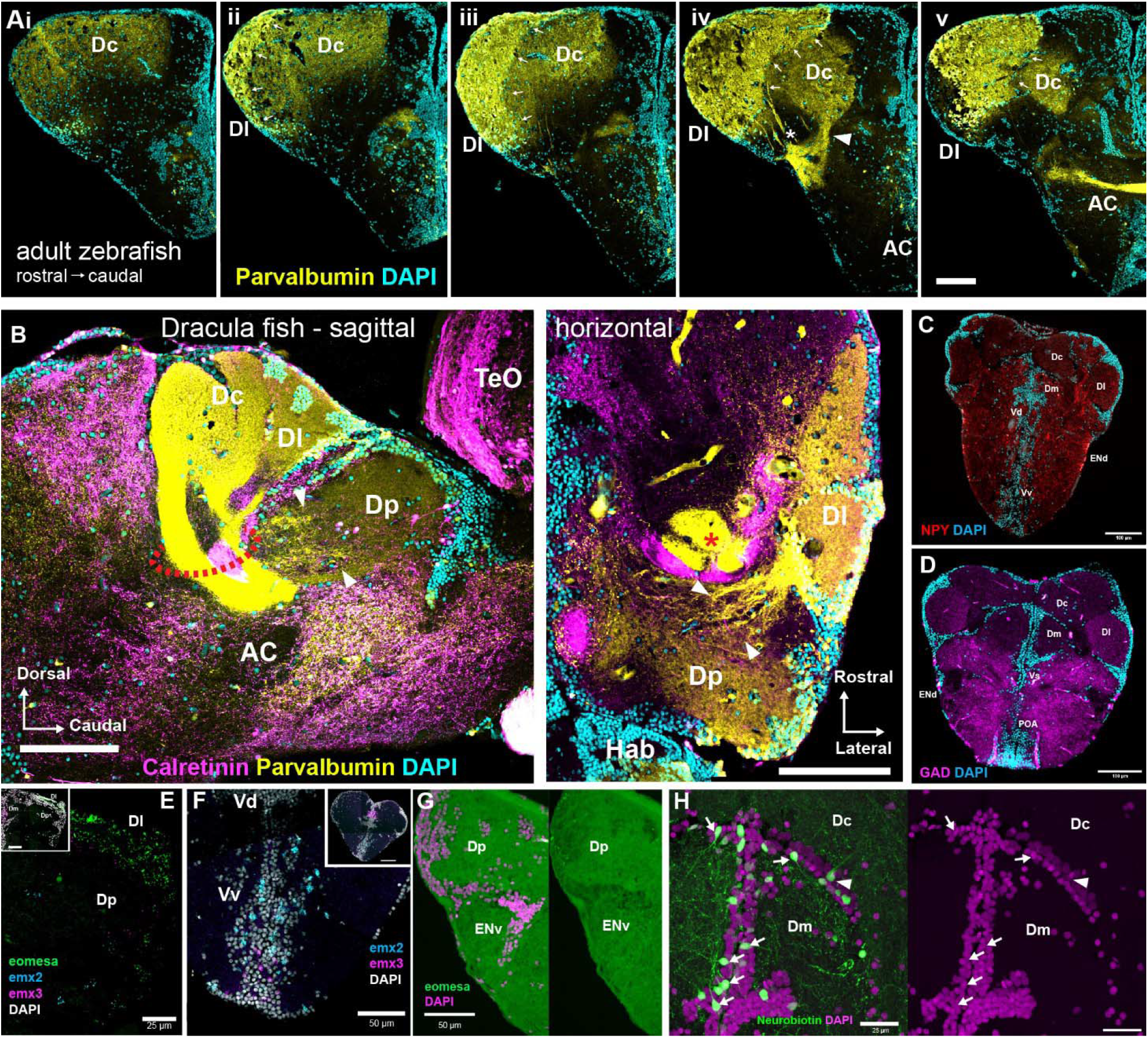
Molecular markers confirm location of caudal pallial divisions in adult Dracula fish (related to Figure 4). (A) Consecutive transverse sections through adult zebrafish telencephalon showing parvalbumin (yellow) immunohistochemistry and DAPI (cyan), progressing from rostral (left) to caudal (right). White arrows denote the border between Dl and Dc. Note that at rostral levels, Dc is at the dorsal surface of the pallium, while caudally it is ventral to Dl. White arrowhead in (iv) points to parvalbumin+ fiber tract connecting Dc to the AC. Asterisk in (iv) denotes parvalbumin+ fibers entering AC that likely emanate from parvalbumin+ cells in Dl. (B) Sagittal and horizontal sections through Dracula fish showing calretinin (magenta) and parvalbumin (yellow) label in caudal pallium. Note parvalbumin staining within the neuropil of Dc and Dl is more intense than in Dp. Red dotted circle in (B) and red asterisk in (C) indicate parvalbumin+ fiber tracts running from Dc and Dl into AC, akin to those shown in zebrafish (Aiv). (C) Tranverse section in Dracula fish shows NPY immunohistochemistry (red) and DAPI (cyan) just rostral to AC. Note NPY staining is lower in Dc as compared to Dm and Dl. (D) Tranverse section shows glutamic acid decarboxylase (GAD, magenta) immunohistochemistry and DAPI (cyan) at the AC. GAD staining is lower in Dc and Dm, and higher in Dl. (E) Caudal telencephalon showing HCR-ISH for *eomesa* (green), *emx2* (cyan) and *emx3* (magenta) in Dp. (F) Rostral telencephalon showing HCR-ISH for *emx2* (cyan) and *emx3* (magenta) localized to Vv. Cell nuclei labeled with DAPI (gray). (G) Light *eomesa* expression (green) in ENv. Cell nuclei labeled with DAPI (magenta). (H) Additional example of neurobiotin filled cells (green) in Dm and Dc. Note cells with smaller somata have processes extending into Dm (arrows) while larger cell has processes extending into Dc (arrowhead).

**Figure S4.**
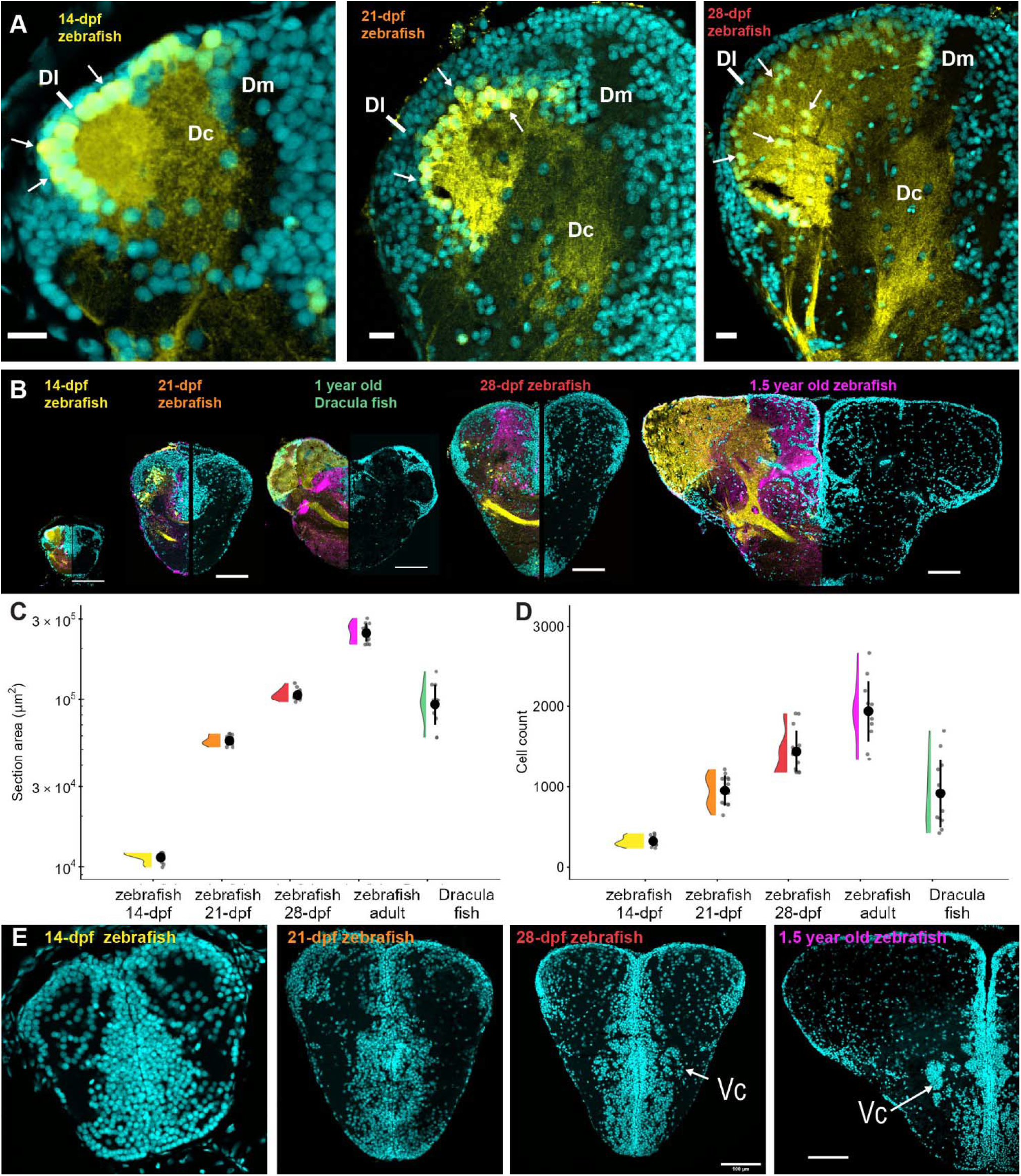
Zebrafish telencephalon across development, compared with adult Dracula fish (related to Figure 5). (A) Zoomed in views of sections from Figure 5A-C showing DAPI (cyan) and parvalbumin+ (yellow) fibers and cells in the caudal pallium of larval zebrafish from 14 to 28-dpf. Parvalbumin+ cells (white arrows) are in the ventricular layer at 14-dpf, are displaced away from it at 21-dpf, and migrate to fill in the interior of Dl by 28-dpf. (B) Comparable transverse sections through caudal telencephalon from 14-, 21-, 28-dpf, 1.5 year old zebrafish and 1 year old Dracula fish, shown at true relative scale. Adult Dracula fish sections fall between 21- and 28-dpf zebrafish. All scale bars = 100 µm. Parvalbumin (yellow), calretinin (magenta), DAPI (cyan). (C) Raincloud plot of telencephalon hemisection area across groups. (D) Raincloud plot of telencephalon hemisection cell count across groups. (E) Transverse DAPI-only images in rostral telencephalon of zebrafish at 14-, 21-, 28-dpf and 1.5 years. Vc is only apparent in 28-dpf and adult zebrafish.

**Figure S5.**
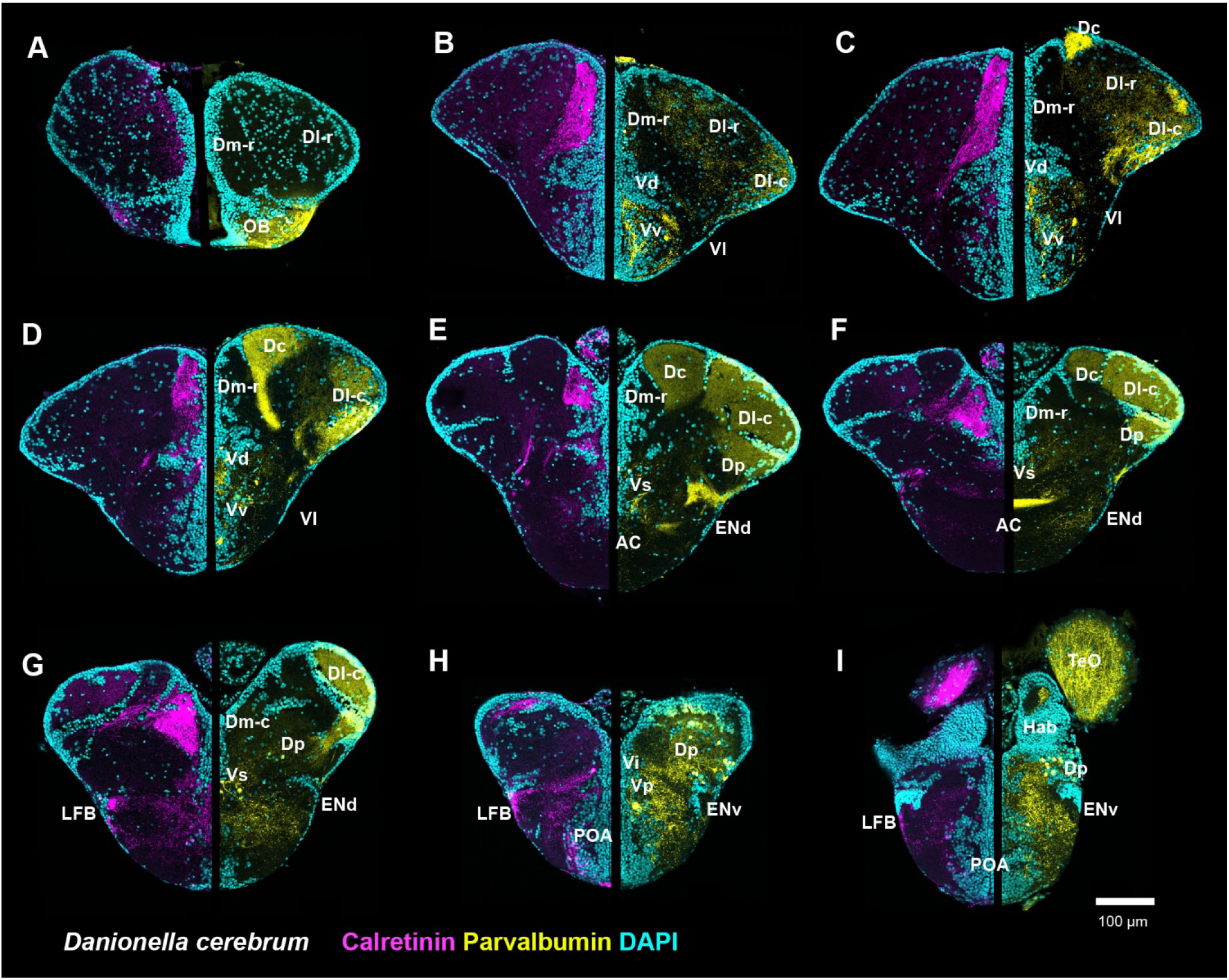
Rostrocaudal patterns of parvalbumin and Calretinin across *Danionella cerebrum* telencephalon (related to Figure 6). (A-I) Transverse sections showing calretinin (magenta) and parvalbumin (yellow) immunohistochemistry with DAPI (cyan) across *D. cerebrum* telencephalon. Most brain regions identified in Dracula fish can be identified in *D. cerebrum* based upon calretinin and parvalbumin labeling along with the comparable cytoarchitecture. The exception is the location of Vi, which is inferred, but requires confirmation with *otpa* expression. See *Telencephalic nomenclature* in Results for telencephalic abbreviations; additional abbreviations: AC = anterior commissure; Hab = habenula, LFB = lateral forebrain bundle; POA = preoptic area; TeO = optic tectum.

**Figure S6.**
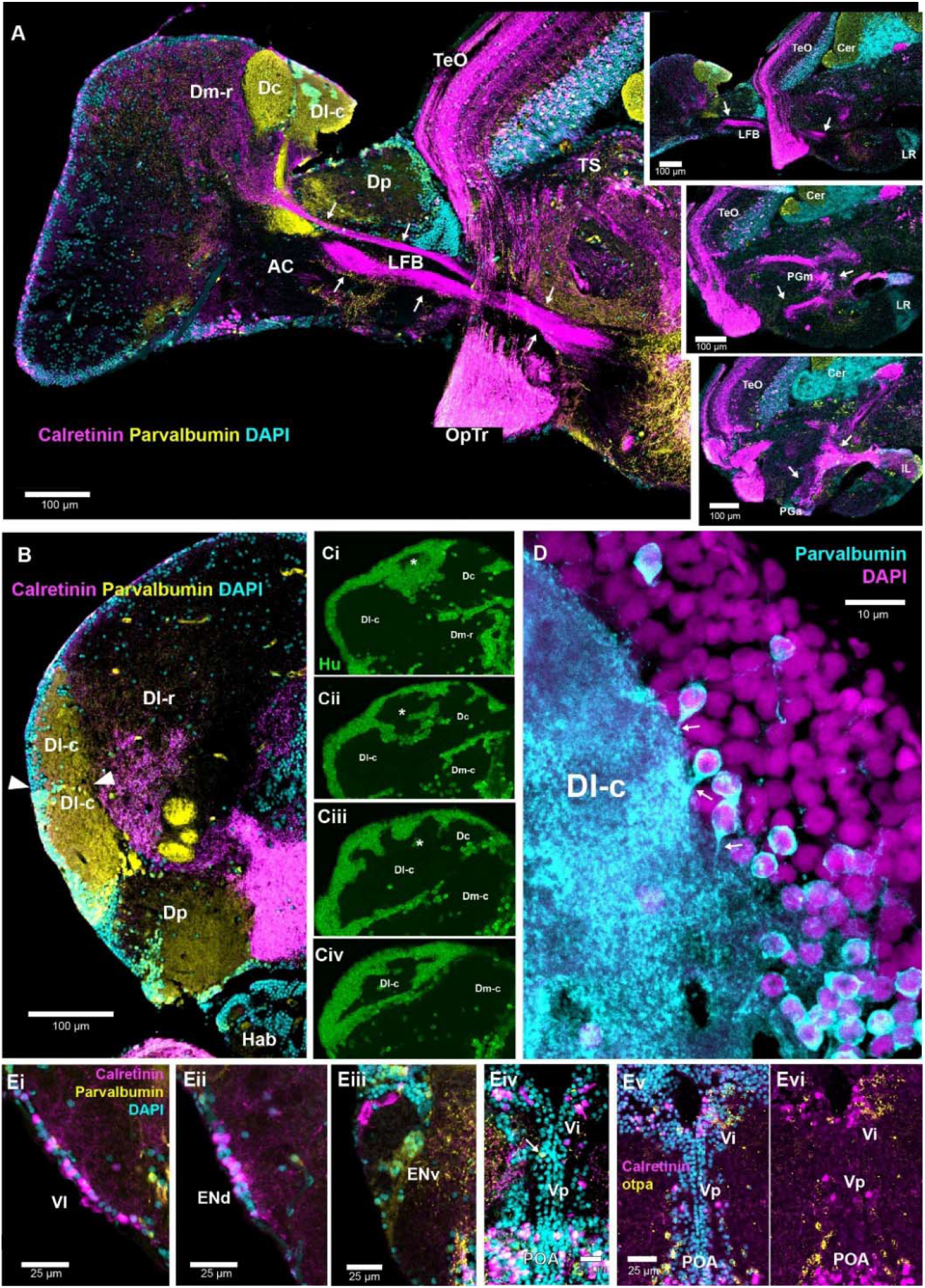
Connectivity and rostrocaudal cytoarchitectural details of the Dracula fish telencephalon (related to Figure 6). (A) Sagittal series showing calretinin+ (magenta) processes in Dm (rostral and caudal, Dm-r, Dm-c) connect via the lateral forebrain bundle (LFB) to the preglomerular complex (anterior and medial, PGa, PGm) in the diencephalon (arrows). Parvalbumin (yellow) and DAPI (cyan). See also Video S1. (B) Horizontal section showing cell band (arrowheads) within the caudal subdivision of Dl (Dl-c). (C) Transverse sections, progressing rostral to caudal, with Hu antibody labeling (green) shows a multi-compartment like structure of Dl-c. Asterisk indicates large cell mass (Ci) that expands (Cii) and then becomes indistinguishable from the rest of Dl-c (Ciii, Civ). (D) High magnification transverse image of parvalbumin somata (cyan) shows processes (arrows) entering the dense parvalbumin neuropil of Dl-c. (E) High magnification transverse images showing calretinin+ somata in Vl (Ei) and ENd (Eii), parvalbumin+ somata in ENv (Eiii). Calretinin+ somata in Vi and Vp, and one parvalbumin+ cell in Vp (Eiv). Colabeling with calretinin+ (magenta) and *otpa* (yellow) confirms calretinin+ somata in Vi and Vp, shown with (Ev) and without (Evi) DAPI (cyan). Abbreviations: AC = anterior commissure; Cer = cerebellum; Hab = habenula; IL = inferior lobe; LR = lateral recess; POA = preoptic area; TeO = optic tectum; TS = torus semicircularis. See *Telencephalic nomenclature* in Results for additional abbreviations.

**Table S1.**
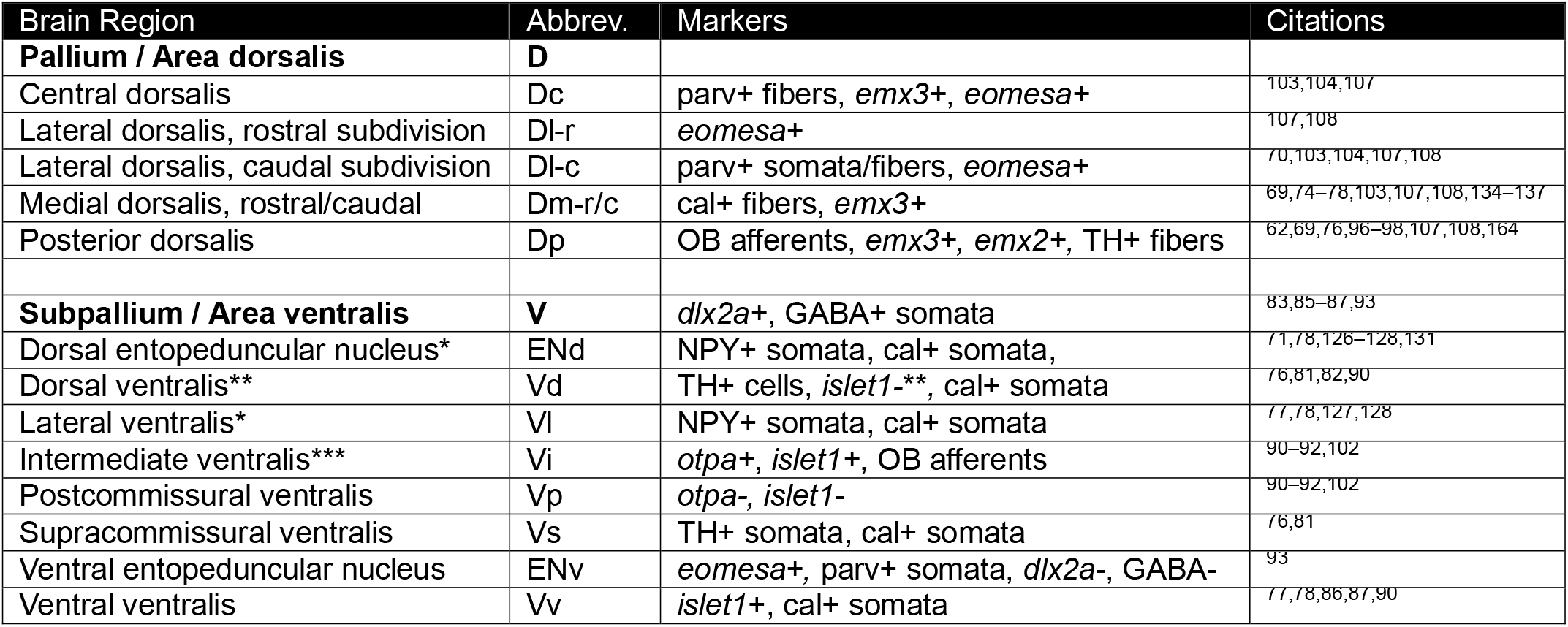
Summary of key markers identifying Dracula fish telencephalic brain regions. Telencephalic brain regions identified in this study and their key diagnostic molecular markers. Citations for each brain region focus on supporting evidence in zebrafish but include other ray-finned fishes if a marker/region relationship is known to be broadly conserved. *ENd and Vl are distinguished from one another by location and morphology of cells. NPY immunohistochemistry shows ENd cells are larger with more prominent projections into pallium. **In zebrafish adults, Vd is largely *islet1* negative, except for a small ventral portion called Vdv^90^. ***The Vi we characterize here follows that of Biechl et al. (2017)^102^, and is likely not equivalent to the Vi of Nieuwenhuys (1963)^45^ and Northcutt and Braford (1980)^62^; described as a lateral extension of Vp. Additional abbreviations: cal = calretinin; NPY = neuropeptide Y, OB = olfactory bulb; parv = parvalbumin. TH = tyrosine hydroxylase

**Table S2.** Species Number, Developmental Stage, Sex and Size. Number of subjects, listed by species, with ratio of males to females (M:F) and standard length (mean ± standard deviation). For zebrafish at 14, 21 and 28-days post fertilization (dpf) are not yet sexually mature. Standard length was measured as distance from snout to caudal peduncle of the tail.^165^ *14-dpf zebrafish larvae have not yet developed a caudal peduncle, so “standard length” was measured as distance from snout to posterior end of the notochord.

| Species, stage | Number | M: F Ratio | Standard Length (SL) |
| --- | --- | --- | --- |
| *Danionella dracula* (Dracula fish), adult | 116 | 1:1 | 1.58 ± 0.14 cm |
| *D. cerebrum*, adult | 6 | 1:1 | 1.28 ± 0.15 cm |
| *D. priapus*, adult | 6 | 1:1 | 2.13 ± 0.14 cm |
| Zebrafish, adult | 6 | 1:1 | 3.06 ± 0.55 cm |
| Zebrafish, 14-dpf | 6 | n/a | 4.82 ± 0.73 mm* |
| Zebrafish, 21-dpf | 6 | n/a | 7.18 ± 0.98 mm |
| Zebrafish, 28-dpf | 6 | n/a | 9.87 ± 1.04 mm |

